# Decoding EGFR ligand bias through an endocytic organelle platform

**DOI:** 10.64898/2026.08.03.742496

**Authors:** Gorana Jendrisek, Deborah Mesa, Stefano Freddi, Giorgia Miloro, Chiara Tordonato, Andrea Francesco Benvenuto, Micaela Quarto, Margherita Caputo, Andrea Raimondi, Giusi Caldieri, Elisa Barbieri, Simone Pelicci, Mario Faretta, Maria Grazia Malabarba, Diego Chianese, Francesca Begnozzi, Paolo Pinton, Massimo Bonora, Pier Paolo Di Fiore, Sara Sigismund

**Author notes:** Correspondence: to SS.

## Abstract

How growth factor receptors decode ligand identity into distinct cellular responses remains a fundamental question in cell signaling. Here, we identify a receptor-proximal mechanism that links ligand-specific EGFR activation to distinct endocytic and biological outputs. We show that EGF, but not TGFα, selectively engages a RAC1-PLCγ2-IP3R signaling axis that supports EGFR non-clathrin endocytosis (NCE). PLCγ2, but not PLCγ1, localizes to RTN3-dependent PM-ER contact sites, where it generates localized Ca²⁺ signals required for completion of NCE, mitochondrial activation and cell motility. This specificity requires the RAC-binding interface of PLCγ2 and is associated with RAC1-dependent formation of CTxB-positive PM regions, indicating that spatial organization contributes to signaling specificity. TGFα fails to efficiently assemble the EGFR-associated organelle platform and instead favors clathrin-dependent EGFR uptake, prolonged proliferative signaling, greater organoid yield, and reduced migration compared with EGF. Together, our findings identify the RAC1-PLCγ2 axis as the key determinant that decodes EGFR ligand bias by coupling receptor trafficking to the metabolic program that supports cell migration.

## Introduction

Growth factor receptor signaling controls multiple cellular outcomes and is finely modulated by different variables, including ligand type, concentration and duration of exposure^1, 2^. The mechanisms through which cells decode this information remain largely unknown.

Our previous work established that epithelial cells sense and decode varying EGF concentrations by engaging distinct endocytic mechanisms that activate specific cellular programs. At low EGF concentrations, EGFR is internalized via CME, which primarily leads to receptor recycling and sustained mitogenic signaling from both the plasma membrane (PM) and endosomal compartments^3^. In contrast, high EGF concentrations activate non-clathrin endocytosis (NCE), which operates in parallel to CME and promotes the assembly of a tripartite organelle platform linking the PM, the endoplasmic reticulum (ER), and mitochondria^4, 5^. These membrane contact sites couple EGFR signaling to mitochondrial metabolism through localized ER Ca^2+^ release that is buffered by adjacent mitochondria, generating Ca^2+^ oscillations and stimulating ATP production at the cell cortex. The resulting increase in local ATP availability drives cortical actin remodeling, which serves a dual purpose: i) it is required for scission of EGFR-NCE tubular invaginations (TIs) and release of NCE carriers into the cytosol, enabling receptor trafficking through the endolysosomal pathway for lysosomal degradation; and ii) it drives cytoskeletal dynamics that support collective cell migration^5^. Thus, ligand concentration determines not only the route of EGFR internalization but also the cellular outcome: at low EGF levels, CME favors receptor recycling and sustained proliferative signaling, whereas at high EGF levels, engagement of NCE redirects activated receptors to lysosomal degradation, thereby limiting the duration and magnitude of mitogenic signaling while promoting migratory responses^3, 5^.

In addition to ligand concentration, EGFR signaling outputs are also influenced by the nature of the activating ligand. Seven different EGFR ligands have been identified, which are associated with distinct cellular responses. A striking example is transforming growth factor alpha (TGFα), which induces a stronger mitogenic response compared with EGF^6^. This differential response has been attributed to differences in receptor conformation upon activation and/or in post-endocytic trafficking, with TGFα primarily directing receptors toward recycling whereas EGF is stronger inducer of lysosomal degradation^7–10^. However, the molecular mechanisms underlying ligand-specific trafficking of the EGFR remain uncharacterized.

In this study, we show that the distinct cellular responses elicited by EGF and TGFα depend on their differential ability to engage the NCE organelle platform. By dissecting the signaling cascade triggered by high EGF concentrations, we identified a RAC1-dependent PLCγ2-IP₃R signaling axis that contributes to internalization, Ca²⁺ oscillations at tripartite contact sites, and changes in mitochondrial membrane potential. PLCγ2 ablation selectively impaired these processes without affecting CME, whereas PLCγ1 had no comparable role. Although both PLCγ paralogs are recruited to activated EGFR, only PLCγ2 localizes to membrane microdomains associated with reticulon 3 (RTN3)-positive ER-PM contact sites, where NCE occurs^4^. Mechanistically, this isoform specificity appears to depend on the RAC-binding interface present in PLCγ2 but not PLCγ1. RAC1 is required for the EGF-induced increase in CTxB-positive PM regions and for ER-PM contact formation. Importantly, this signaling axis is preferentially engaged by EGF compared with TGFα, thereby conferring ligand-specific EGFR responses. Consistent with this, TGFα yielded more organoids above prespecified size thresholds than EGF, a phenotype recapitulated in NCE-defective RTN3-KO organoids grown in EGF, indicating that NCE activation restrains EGF-dependent mitogenic signaling. Conversely, inefficient engagement of this pathway by TGFα is associated with shorter-lived Ca²⁺ oscillations, no detectable increase in mitochondrial membrane potential, and reduced migratory capacity compared with EGF.

Together, our findings identify a receptor-proximal signaling axis that couples the EGFR-NCE organelle platform with specific biological outputs, providing a mechanistic explanation for how ligand identity is translated into distinct biological responses and for the long-standing observation that TGFα is a more potent mitogen than EGF.

## Results

### PLCγ2 is a critical component of the EGFR-NCE pathway

To identify signaling molecules specifically associated with EGFR-NCE, we investigated the activation of major EGFR downstream effectors (AKT, ERK1/2, SHC, PLCγ) following stimulation of HeLa cells with either low (1 ng/ml) or high (100 ng/ml) concentrations of EGF. Only the PLCγ family members (PLCγ1 and PLCγ2) displayed selective phosphorylation in response to high-dose EGF over a range of stimulation times, whereas low-dose EGF failed to induce their activation (**Fig. 1A**). Consistent with this observation, dose-response analysis at 2 min revealed a sharp increase in PLCγ phosphorylation between 3 and 10 ng/ml EGF, reaching maximal levels at 30 ng/ml (**Fig. S1A**). Notably, this activation profile closely mirrored the dose dependence of EGFR-NCE, which is activated above 10 ng/ml EGF, reaching a maximum at 30 ng/ml^11^. In contrast, SHC phosphorylation increased gradually with increasing EGF doses (**Fig. S1A**). These findings indicate that PLCγ activation closely correlates with engagement of the EGFR-NCE pathway. To determine whether PLCγ enzymes have a functional role in EGFR-NCE, we assessed the effects of their knockdown (KD) on the internalization of the NCE-specific cargo, CD147^4^. Confocal microscopy of cells stimulated with Alexa Fluor 555-labeled EGF at high concentration revealed that depletion of PLCγ2, but not PLCγ1, inhibited the internalization of both EGF and CD147 (**Fig. 1B**). This inhibitory effect of PLCγ2 KD was not enhanced by combined PLCγ1/2 KD, indicating that PLCγ1 does not function redundantly with PLCγ2 in this pathway (**Fig. 1B**, right panel, and **S1B**). The requirement for PLCγ2 was confirmed using independent RNAi oligonucleotides (**Fig. S1C,D**, see also **Fig. S1E** for an additional PLCγ1 oligonucleotide). Moreover, the inhibitory effect of PLCγ2 KD was quantitatively comparable to that observed upon KD of the established NCE regulator RTN3, and combined RTN3/PLCγ2 depletion did not produce an additive effect, consistent with both proteins acting within the same pathway (**Fig. S1F**).

**Figure 1.**
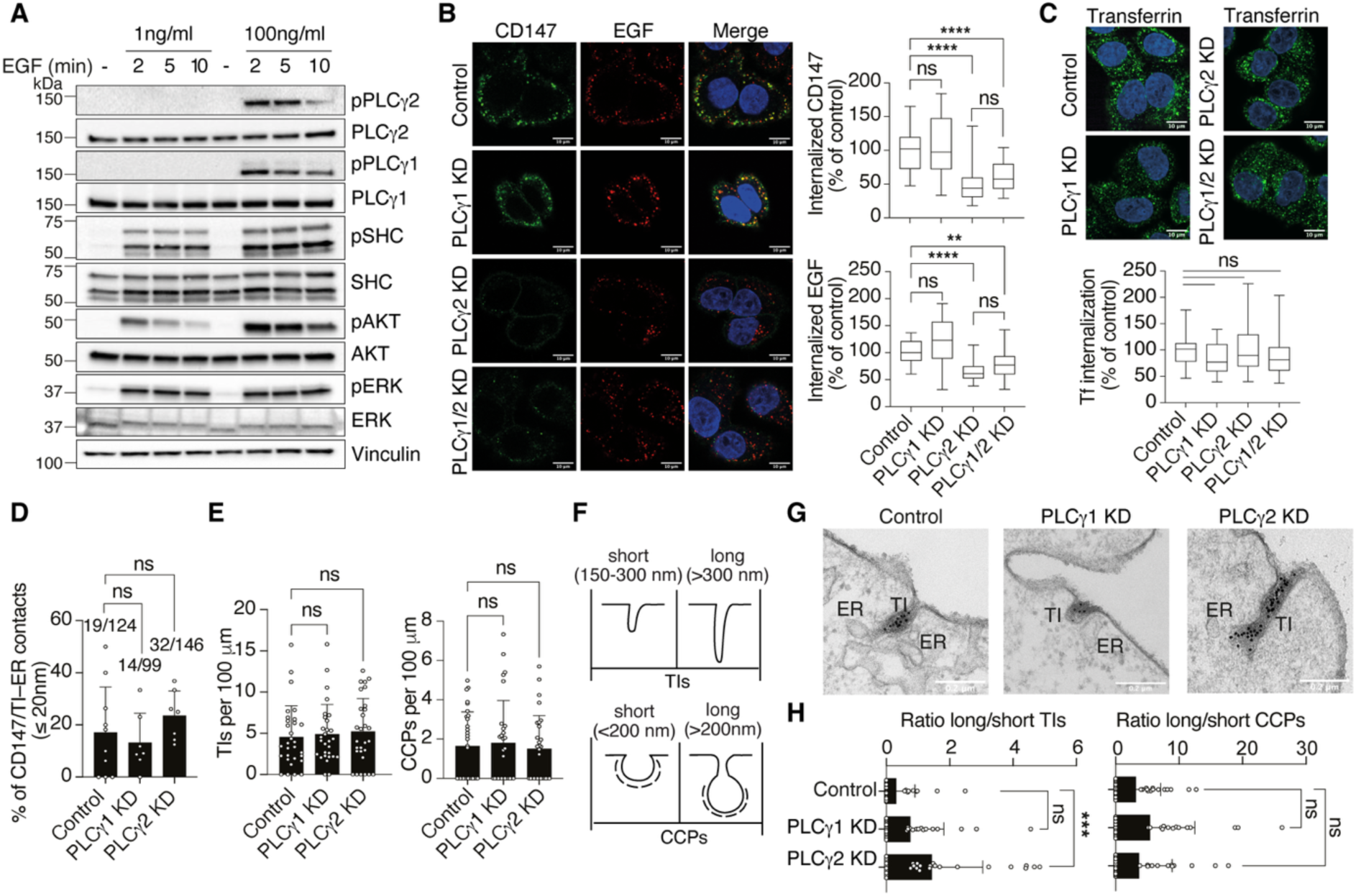
PLCγ2, but not PLCγ1, is required for EGFR-NCE. **A.** PLCγ enzymes are selectively activated by high-dose EGF. HeLa cells were serum-starved for 16 h (−), stimulated with low (1 ng/ml) or high (100 ng/ml) EGF, and harvested for immunoblot analysis at the indicated times. Vinculin, loading control. MW markers shown on the left. **B.** PLCγ2 depletion inhibits EGFR-NCE. CD147 internalization was assessed by IF in HeLa cells following the indicated KDs or mock treatment and stimulation with high-dose Alexa Fluor-555 EGF (∼40 ng/ml) for 8 min. Left, representative confocal images: CD147, green; EGF, red; blue, DAPI. Scale bar, 10 µm. Right, quantification of internalized CD147 (upper panel) and EGF (lower panel), expressed as percentage of control (box plots of mean integrated fluorescence intensity). CD147: N (fields), Control = 28, PLCγ1 KD = 28, PLCγ2 KD = 28, PLCγ1/2 KD = 30; n = 3. EGF: N (fields), Control = 29, PLCγ1 KD = 27, PLCγ2 KD = 29, PLCγ1/2 KD = 31; n = 3. One-way ANOVA: **P < 0.01; ****P < 0.0001; ns. **C.** CME is unaffected by PLCγ depletion. HeLa cells as in B were monitored for Alexa Fluor-488 transferrin (Tf; 50 μg/ml) internalization. Upper, representative confocal images: Tf, green; blue, DAPI. Scale bar, 10 μm. Lower, quantification of internalized Tf, expressed as percentage of control (box plots of mean integrated fluorescence intensity). N (fields): Control = 32, PLCγ1 KD = 32, PLCγ2 KD = 32, PLCγ1/2 KD = 32; n = 3. One-way ANOVA: ns. **D.** PLCγ depletion does not affect ER recruitment to CD147-positive endocytic structures. ER proximity of CD147-positive tubular invaginations (TIs) was assessed by EM morphometry in HeLa cells treated with the indicated KDs and stimulated with high-dose EGF (30 ng/ml) for 5 min. The percentage of structures in contact with the ER (≤ 20 nm) is shown as mean ± SD. N (cells): Control = 10, PLCγ1 KD = 7, PLCγ2 KD = 7; n = 1. One-way ANOVA: ns. **E.** PLCγ depletion does not affect the abundance of EGFR endocytic intermediates. Quantification of NCE TIs and CCPs in HeLa cells treated as in “D”, analyzed by EM using gold-labeled EGFR. The number of gold-labeled structures per 100 μm of PM is shown as mean ± SD. N (100 μm PM lengths): TIs, Control = 30, PLCγ1 KD = 29, PLCγ2 KD = 29; CCPs, Control = 30, PLCγ1 KD = 29, PLCγ2 KD = 29; n = 1. One-way ANOVA: ns. **F-H.** PLCγ depletion causes accumulation of elongated EGFR-NCE-TIs. EM analysis of EGFR-containing endocytic intermediates in HeLa cells treated as in D. F, schematic defining short and long TIs and CCPs. G, representative EM images of EGFR-positive endocytic structures. Scale bar, 200 nm. H, ratio of long to short TI or CCP structures per 100 μm of PM, shown as mean ± SD. N (100 μm PM lengths): Control = 30, PLCγ1 KD = 29, PLCγ2 KD = 29. One-way ANOVA: ***P < 0.001, ns.

Notably, the role of PLCγ2 was specific to EGFR-NCE, as depletion of either PLCγ2 or PLCγ1 had no effect on CME, as assessed by internalization of the canonical CME cargo transferrin (Tf) (**Fig. 1C**). Moreover, the selective requirement for PLCγ2, but not PLCγ1, in EGFR-NCE was confirmed in the normal human keratinocyte cell line HaCaT (**Fig. S2**), in which EGFR-NCE has been shown to be activated^4, 5^.

To determine at which stage of the EGFR-NCE pathway PLCγ2 functions, we performed EM morphometry. Our previous work established that EGFR-NCE proceeds through three sequential stages: 1) ER-PM contact site formation requiring RTN3; 2) TI formation and elongation requiring ER-PM-mitochondria contact sites; and 3) TI fission requiring dynamin, actin, Ca^2+^ and ATP^5^. We used gold-labeled EGFR to visualize EGFR-containing endocytic intermediates in HeLa cells stimulated with high-dose EGF for 5 min. Neither PLCγ1 or PLCγ2 KD affected the formation of ER-PM contact sites (**Fig. 1D**) or the total number of EGFR-positive TIs (**Fig. 1E**). However, PLCγ2 depletion significantly increased the ratio of long (>300 nm) to short (150-300 nm) TIs, whereas PLCγ1 KD had a small, nonsignificant effect (**Fig. 1G,H**). The effects of PLCγ2 KD were comparable to those observed upon inhibition of either the IP3R or dynamin^4, 5^, consistent with a role for PLCγ2 in TI fission. In contrast, neither PLCγ1 nor PLCγ2 depletion altered the number or length of EGFR-containing clathrin-coated pits (CCPs) (**Fig. 1E**, right, and **Fig. 1H, right**), further supporting a selective role of PLCγ2 in EGFR-NCE. Together, these results indicate that PLCγ2 is dispensable for ER-PM contact site formation but is required for efficient fission of EGFR-NCE TIs.

### PLCγ2 couples EGFR activation to NCE-associated Ca²⁺ signaling and changes in mitochondrial membrane potential

Growth factor stimulation typically induces the recruitment of PLCγ enzymes to receptors at the PM, where they are activated and catalyze the hydrolysis of phosphatidylinositol 4,5-bisphosphate (PIP₂), generating diacylglycerol (DAG) and IP3. IP3 subsequently activates IP3R channels at the ER, triggering Ca^2+^ release into the cytosol (**Fig. 2A**).

**Figure 2.**
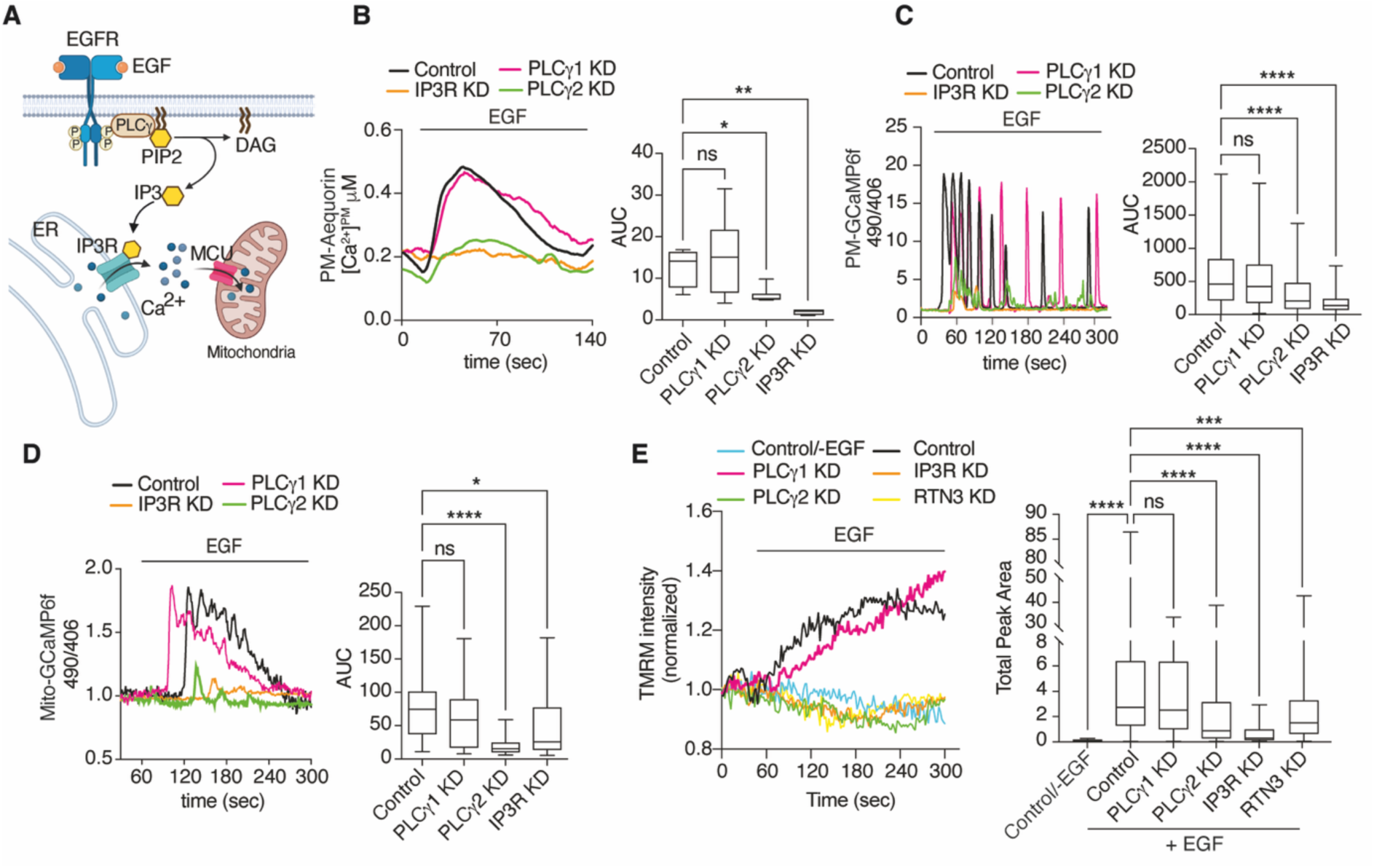
PLCγ2 couples EGF activation to Ca^2+^ signaling and mitochondrial bioenergetics. **A.** Schematic of PLCγ activation downstream of EGF stimulation. EGFR activation recruits PLCγ to the PM, where it is phosphorylated and hydrolyzes PIP2 into DAG and IP3. IP3 activates IP3R on the ER, triggering Ca^2+^ release from the ER lumen. Created with BioRender.com. **B, C**. PLCγ2 depletion abolishes EGF-induced PM-localized Ca²⁺ signaling. HeLa cells expressing PM-targeted Ca^2+^ sensors, PM-Aequorin (B) or PM-GCaMP6f (C), were subjected to the indicated KDs and stimulated with high-dose EGF (100 ng/ml). Left, representative Ca²⁺ traces showing population responses measured with PM-Aequorin or single-cell responses measured with PM-GCaMP6f. For PM-GCaMP6f, the 490/406 nm emission ratio is shown. Right, box plots of Ca^2+^ response AUC. Panel B: N (coverslips, whole cell population): Control = 12 (n = 3), PLCγ1 KD = 10 (n = 3), PLCγ2 KD = 8 (n = 3), IP3R KD = 4 (n = 1). Panel C: N (cells): Control = 191, PLCγ1 KD = 161, PLCγ2 KD = 202, IP3R KD = 173; n = 3. One-way ANOVA: *P < 0.05; **P < 0.01; ****P < 0.0001; ns. **D.** PLCγ2 depletion inhibits EGF-induced mitochondrial Ca²⁺ uptake. HeLa cells expressing the mitochondria-targeted Ca^2+^ sensor, Mito-GCaMP6f, were assessed using the experimental conditions and statistical analysis as in C. N (cells): Control = 27, PLCγ1 KD = 28, PLCγ2 KD = 32, IP3R KD = 22; n = 3. **E.** PLCγ2 depletion impairs EGF-induced activation of mitochondrial bioenergetics. Mitochondrial membrane potential was monitored by TMRM fluorescence in HeLa cells treated with KDs and stimulated with high-dose EGF (100 ng/ml). Mock KD and serum-starved (−EGF) controls were included. Left, representative time course of TMRM fluorescence. Right, box plots of AUC. N (cells): -EGF Control = 51, Mock Control = 492, PLCγ1 KD = 444, PLCγ2 KD = 480, IP3R KD = 73, RTN3 KD = 439; n = 3 (except for -EGF Control and IP3R KD; n = 1). One-way ANOVA: ***P < 0.001; ****P < 0.0001; ns.

To determine whether high-dose EGF activates this canonical PLC**γ−**IP3R signaling cascade during EGFR-NCE, we monitored subplasmalemmal Ca²⁺ dynamics using two PM-targeted Ca^2+^ sensors: PM-Aequorin, which measures Ca²⁺ responses at the cell population level, and PM-GCaMP6f, which enables single-cell monitoring. High-dose EGF induced robust Ca²⁺ signals in HeLa cells, detected either as a transient wave with PM-Aequorin or as repetitive oscillatory spikes with PM-GCaMP6f (**Fig. 2B,C**). These responses were markedly reduced following depletion of PLCγ2 or IP3R, whereas PLCγ1 KD had no significant effect (**Fig. 2B,C**). Thus, the localized EGF-induced Ca²⁺ response at the PM depends on the PLCγ2-IP₃R axis.

Our previous work demonstrated that these Ca²⁺ signals are generated at PM-ER-mitochondria contact sites, where mitochondrial Ca²⁺ uptake is required to sustain Ca²⁺ oscillations (**Fig. 2A**)^5^. Consistent with this model, measurements with the mitochondrial matrix-targeted sensor Mito-GCaMP6 revealed rapid mitochondrial Ca²⁺ uptake following high-dose EGF stimulation. Importantly, mitochondrial Ca²⁺ signals were abolished by PLCγ2 or IP₃R depletion but were unaffected by PLCγ1 KD (**Fig. 2D**).

Given that mitochondrial Ca²⁺ uptake stimulates oxidative metabolism, we next examined mitochondrial activation by measuring changes in mitochondrial membrane potential. High-dose EGF induced a rapid increase in TMRM fluorescence, consistent with mitochondrial hyperpolarization (**Fig. 2E**). This response was inhibited by depletion of PLCγ2 or IP3R, but not PLCγ1 (**Fig. 2E**).

Together, these findings identify PLCγ2 as the upstream activator of the localized Ca^2+^ signaling and mitochondrial membrane-potential response previously shown to drive ATP production at PM-ER-mitochondria contact sites and promote fission of EGFR-NCE TIs^5^.

### EGF selectively recruits PLCγ2 to RTN3-positive EGFR-NCE sites at the PM

Although both PLCγ family members are efficiently activated by high-dose EGF, as evidenced by their phosphorylation (**Fig. 1A** and **Fig. S1A**), only PLCγ2 is required for EGFR-NCE. We therefore hypothesized that the functional specificity of PLCγ2 might arise from its selective recruitment to NCE sites at the PM. Consistent with this possibility, PLCγ2, but not PLCγ1, has been reported to localize to cholesterol- and sphingolipid-rich membrane domains in activated lymphocytes^12^.

To test this hypothesis, we generated stable HeLa cell lines expressing comparable levels of HA-tagged PLCγ2 or PLCγ1 (**Fig. S3A**). EGFR activation in both cell lines was indistinguishable from that observed in empty vector control cells (**Fig. S3B,C**). Under basal conditions, both PLCγ enzymes localized mainly in the cytoplasm, whereas after 2 min of high-dose EGF stimulation, a fraction of both PLCγ2 and PLCγ1 relocalized to the PM (**Fig. 3A**). STORM super-resolution microscopy confirmed the recruitment of both enzymes to PM regions enriched in EGFR clusters (**Fig. 3B, Fig. S4A**), consistent with their EGF-induced phosphorylation (**Fig. 1A**).

**Figure 3.**
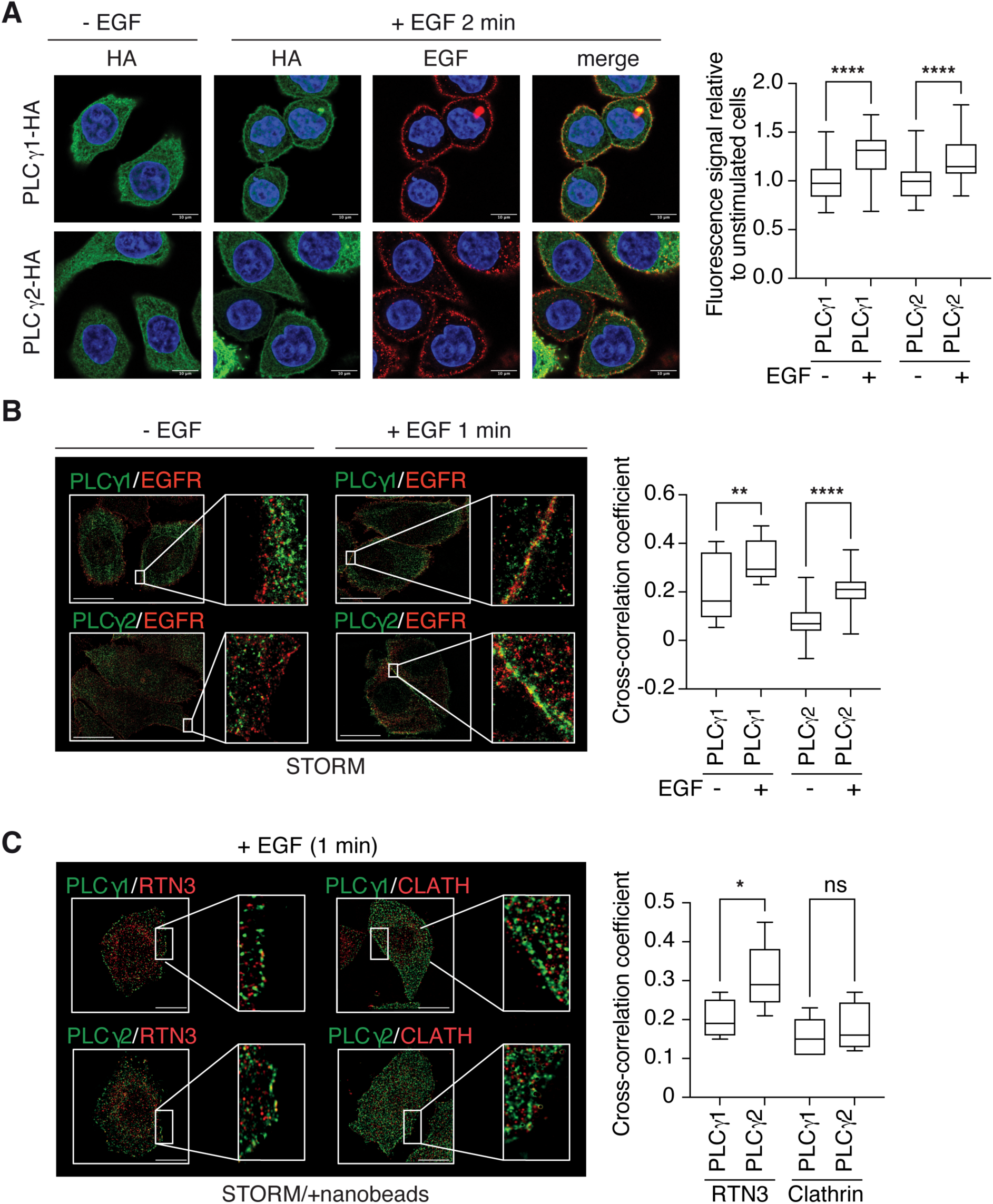
PLCγ2 is selectively recruited to RTN3-positive EGFR-NCE sites at the PM. **A.** PLCγ enzymes are recruited to the PM following EGF stimulation. HeLa cells stably expressing HA-tagged PLCγ1 or PLCγ2 were serum starved for 2h, then stimulated with high-dose Alexa Fluor-555 EGF (∼40 ng/ml) for 2 min or left untreated (–EGF) and analyzed by IF confocal microscopy. Left, representative images: EGF, red, PLCγ-HA; green, DAPI, blue. Scale bar, 10 μm. Right, quantification of PLCγ-HA recruitment to the PM, measured as fluorescence intensity within 0.5 µm of the cell periphery and normalized to unstimulated cells. N (cells): -EGF, PLCγ1 = 40, PLCγ2 = 58; +EGF, PLCγ1 = 40, PLCγ2 = 45. One-way ANOVA: ****P < 0.0001. **B.** EGF induces co-clustering of PLCγ enzymes and EGFR at the PM. HeLa cells as in A were stimulated with high-dose EGF (100 ng/ml) for 1 min and analyzed by IF STORM microscopy. Left, representative STORM images (median optical section): EGFR, red; PLCγ-HA, green. Scale bar, 15 μm. Right, quantification of EGFR and PLCγ-HA co-clustering within 40 nm of the cell periphery by Spearman cross-correlation analysis. N (cells): -EGF: PLCγ1 = 12, PLCγ2 = 24; +EGF: PLCγ1 = 11, PLCγ2 = 26. A representative experiment is shown (n = 2). One-way ANOVA: **P < 0.01; ****P < 0.0001. **C.** PLCγ2 but not PLCγ1 specifically co-clusters with RTN3 in the vicinity of the PM in the presence of high EGF. HeLa cells as in A were stimulated with high-dose EGF (100 ng/ml) for 1 min and analyzed by STORM microscopy. Dark areas correspond to the nanobeads used as fiducial markers for drift correction and two-channel alignment (see **Fig. S4C**). After channel registration, bead-containing regions were masked in the images used for figure preparation. Left, representative STORM images (median optical section): RTN3/Clath, red; PLCγ-HA, green. Scale bar, 10 μm. Right, quantification of co-clustering as in B. N (cells): RTN3 PLCγ1 = 4, RTN3 PLCγ2 = 5, Clath PLCγ1 = 5, Clath PLCγ2 = 4. A representative experiment is shown (n = 2). One-way ANOVA: *P < 0.05, ns.

Despite this common recruitment to activated EGFR, PLCγ2 exhibited greater spatial correlation with the NCE regulator RTN3 at the cell periphery than did PLCγ1 (**Fig. 3C**, controls in **Fig. S4B**). In contrast, PLCγ1 showed little association with RTN3 despite its efficient recruitment to the PM. No difference in co-clustering with clathrin at the PM (**Fig. 3C**) or with RTN3 in the cytosol (**Fig. S4C**) between PLCγ1 and PLCγ2 is observed. These findings suggest that, although both PLCγ enzymes are recruited to activated EGFR, PLCγ2 is preferentially enriched at RTN3-positive NCE sites, providing a spatial basis for its specific role in EGFR-NCE.

### RAC1 is required for EGF-induced PM microdomain assembly and ER-PM contact formation during EGFR-NCE

The selective recruitment of PLCγ2 to EGFR-NCE sites may reflect its unique ability to interact with the small Rho GTPase RAC through its split PH domain, and to be activated by this interaction^13, 14^. RAC itself is activated downstream of EGFR through the PI3K-PIP3 pathway and the RAC GEF, TIAM^15, 16^, accumulates in cholesterol-enriched PM domains^17^ and contributes to EGF-induced cortical actin remodeling and downstream EGFR signaling ^18^. To investigate the involvement of RAC proteins in EGFR-NCE, we first examined their expression in HeLa cells and found that RAC1, but not RAC2 or RAC3, was detectably expressed (**Fig. S5A**). RAC1 depletion markedly impaired internalization of both the NCE cargo CD147 and EGFR following high-dose EGF stimulation (**Fig. 4A**) and prevented the formation of ER-PM contact sites (**Fig. 4C,D**). Notably, RAC1 exerts a broader role in endocytosis, as RAC1 depletion also reduced internalization of the CME cargo Tf (**Fig. S5B**). EGFR-NCE depends on PM microdomains enriched in cholesterol and glycosphingolipids^5, 19^. Consistent with this, high-dose EGF markedly increased PM staining by cholera toxin B (CTxB), a probe for the glycosphingolipid GM1 (**Fig. 4B**)^20^. RAC1 knockdown abolished this increase, suggesting that RAC1 affects the abundance or organization of CTxB-accessible GM1 at the PM.

**Figure 4.**
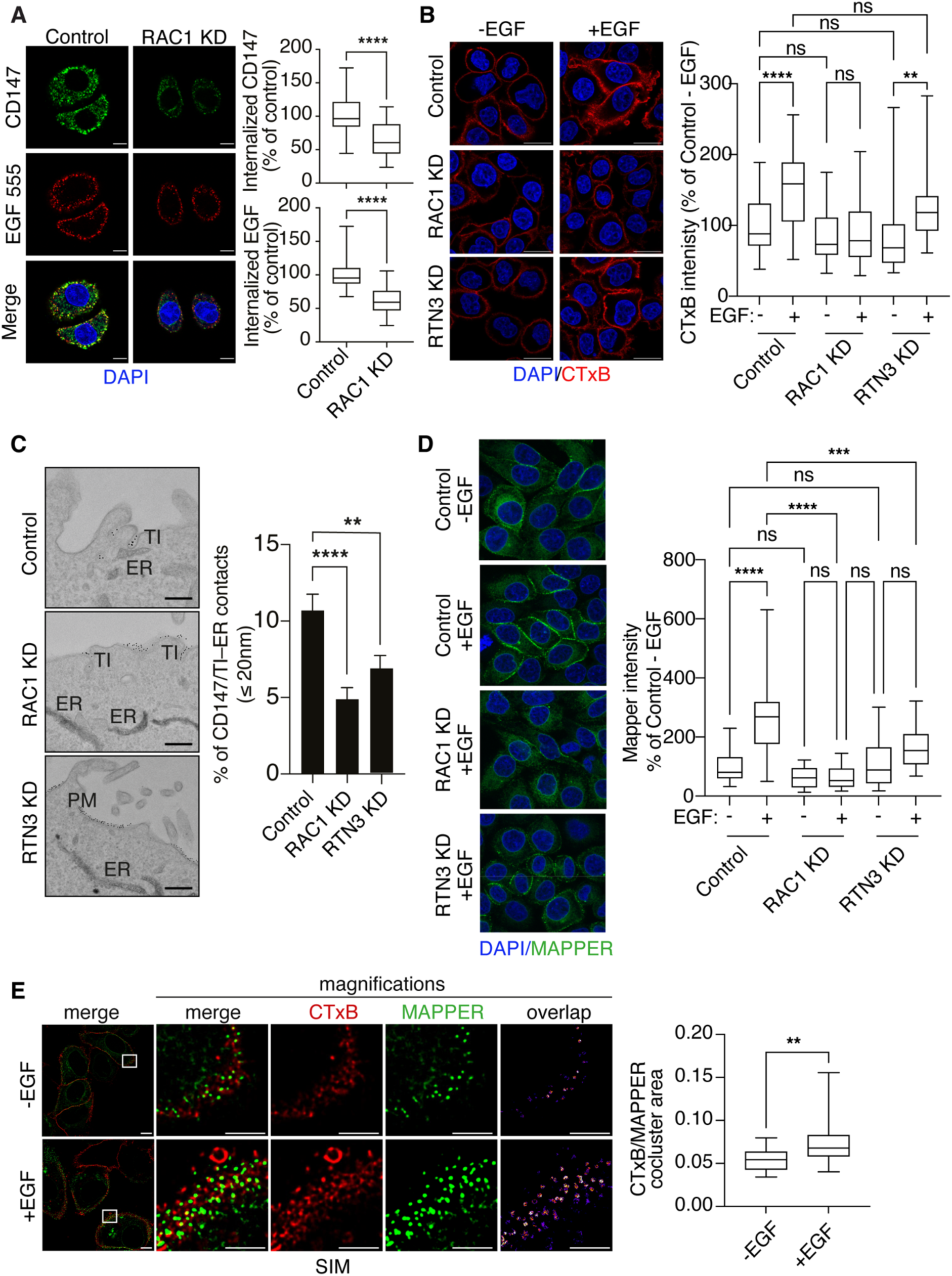
RAC1 drives formation of GM1-rich PM microdomains and ER-PM contact sites. **A.** RAC1 depletion inhibits EGFR-NCE. CD147 internalization was monitored by IF in HeLa cells subjected to RAC1 KD or mock treatment and stimulated with high-dose Alexa Fluor-555 EGF (∼40 ng/ml, red) for 5 min. Left, representative confocal images: CD147, green; EGF, red; DAPI, blue. Scale bar, 10 µm. Right, quantification of internalized CD147 (upper) and EGF (lower) expressed as percentage of control (box plot of mean integrated fluorescence intensity). CD147, N (fields): Control = 28, RAC1 KD = 26; n = 3. EGF, N (fields): Control = 30, RAC1 KD = 29; n = 3. Student’s two-tailed t-test; ****P < 0.0001. **B.** RAC1, but not RTN3, is required for formation of GM1-rich PM microdomains. HeLa cells were subjected to RAC1, RTN3 or mock KD, serum starved for 2 h, then stimulated (+EGF) or not (-EGF) with high-dose EGF (100 ng/ml) for 1 min. Cells were fixed, incubated with Alexa Fluor 555-conjugated cholera toxin B subunit (CTxB, 285 ng/ml), permeabilized and analyzed by confocal microscopy. Left, representative IF images: CTxB, red; DAPI, blue. Scale bar, 10 µm. Right, quantification of CTxB binding as percentage of unstimulated control (box plot of mean integrated fluorescence intensity). N (fields): -EGF, Control = 30, RAC1 KD = 26, RTN3 KD = 20. N (fields): +EGF, Control = 29, RAC1 KD = 31, RTN3 KD = 20. n = 3 except for RTN3 KD (n = 2). One-way ANOVA: **P < 0.01; ****P < 0.0001; ns. **C.** RAC1 depletion inhibits EGF-induced ER recruitment to CD147-positive endocytic structures. ER proximity of CD147-positive endocytic structures was assessed by EM morphometry using gold-labeled CD147 in HeLa cells subjected to the indicated KDs and stimulated with high-dose EGF (30 ng/ml) for 5 min. Left, representative EM images. Scale bar, 500 nm. Right, percentage of CD147-positive structures in contact with the ER (≤ 20 nm) shown as mean ± SEM. N (cells): Control = 20, RAC1 KD = 20, RTN3 KD = 20. One-way ANOVA: ****P < 0.0001, **P < 0.01. **D.** RAC1 depletion abolishes formation of EGF-induced ER-PM contact sites. HeLa cells expressing pSlik-GFP-MAPPER were subjected to indicated KDs, GFP-MAPPER protein expression was induced with 1 μg/ml of doxycycline overnight (16 hours). Cells were stimulated with high dose Alexa Fluor-555 EGF (∼40ng/ml) for 5 min. Left, representative confocal images. Right, quantification of MAPPER intensity normalized on % of Control – EGF (box plot). Scale bar, 10 µm. N (fields): -EGF, Control = 17, RAC1 KD = 18, RTN3 KD = 19; +EGF, Control = 20, RAC1 KD = 18, RTN3 KD = 20 (n = 2). One-way ANOVA: **P < 0.01; ****P < 0.0001; ns**. E.** EGF induces co-clustering of MAPPER-positive ER-PM contacts and CTxB-positive PM microdomains. HeLa cells expressing pSlik-GFP-MAPPER were serum starved for 6 h, stimulated with high-dose EGF (100 ng/mL) for 1 min, or left unstimulated. Cells were fixed and then incubated with CTxB (285 ng/ml). Left, representative structured illumination microscopy (SIM) images of CTxB–MAPPER co-clusters in -/+ EGF conditions: Mapper (green); CTxB (red). Scale bar, 5 μm. Right, quantification of CTxB– MAPPER co-cluster area. N (fields of view): -EGF = 33, +EGF = 23 (n = 1). One-way ANOVA: **P < 0.01.

Because EGFR-NCE is initiated at RTN3-dependent ER-PM contact sites, we next investigated whether RAC1 also regulates their formation. By EM, using CD147-gold to label NCE tubular invaginations and HRP-KDEL to visualize the ER, RAC1 depletion markedly reduced the frequency of ER-PM contacts, to an extent comparable to the RTN3 KD positive control (**Fig. 4C**). This result was independently confirmed using the fluorescent ER-PM contact site reporter MAPPER^21^. Whereas EGF stimulation induced a robust increase in MAPPER-positive puncta, this response was almost completely abolished in RAC1-depleted cells (**Fig. 4D**).

Finally, CTxB-positive membrane domains showed co-clustering with MAPPER-positive ER-PM contact sites, and this association increased upon EGF stimulation (**Fig. 4E**). These observations are consistent with the notion that EGF-induced ER-PM contact sites are assembled within CTxB-positive PM regions where EGFR-NCE is initiated.

Consistent with a model in which RAC1 contributes to preferential PLCγ2 activation during EGFR-NCE, RAC1 KD markedly reduced EGF-induced PLCγ2 phosphorylation, whereas it had only a modest effect on PLCγ1 phosphorylation (**Fig. 5A**). Consequently, the localized Ca²⁺ response at the PM was strongly impaired (**Fig. 5B**), to an extent comparable to PLCγ2 ablation. Together, these findings are consistent with RAC1 acting upstream of PLCγ2 activation during EGFR-NCE and suggest a model in which RAC1 promotes the selective activation of PLCγ2 within NCE-associated PM microdomains.

**Figure 5.**
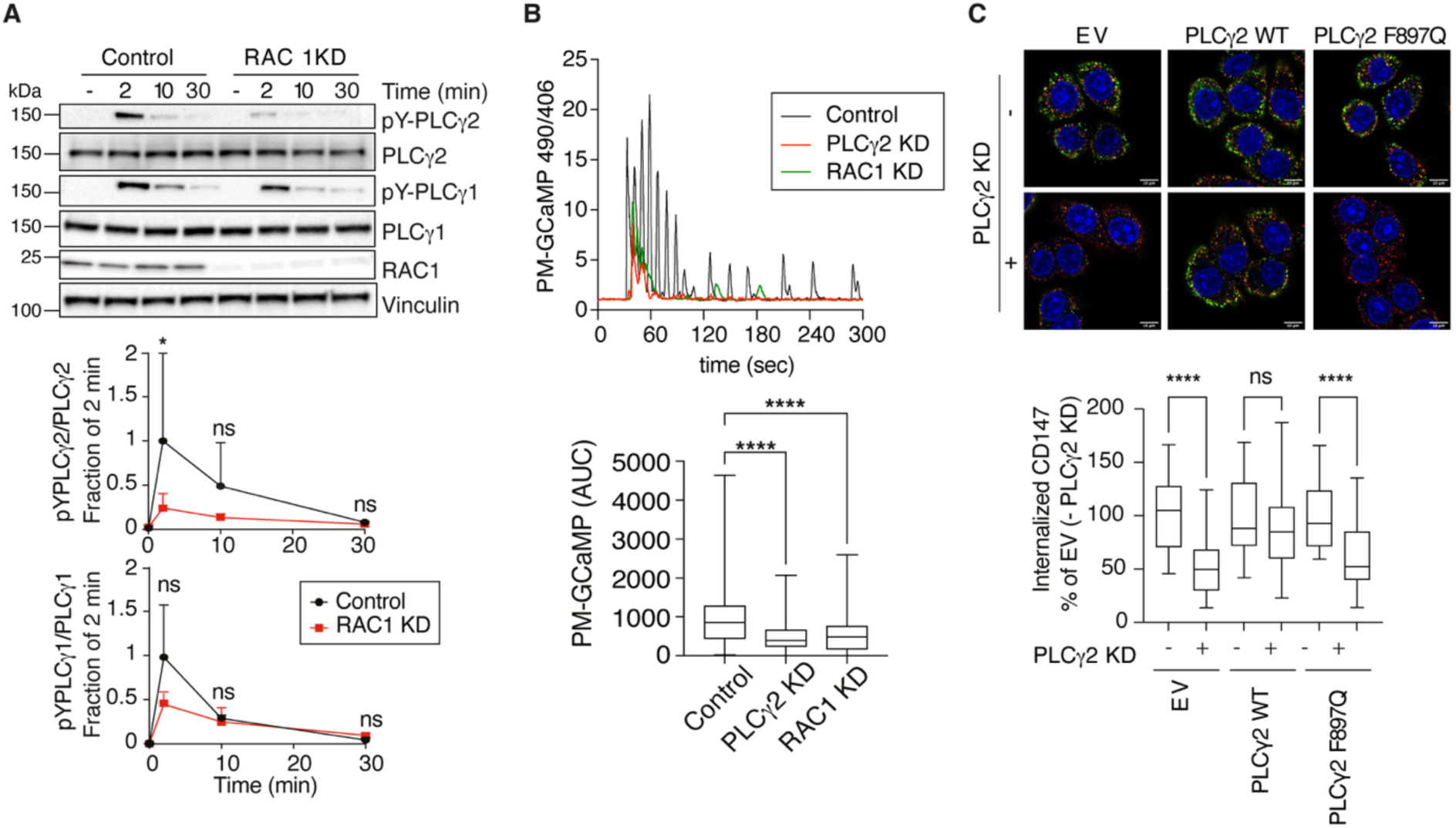
RAC1 activates PLCγ2 to promote EGFR-NCE. **A.** RAC1 acts upstream of PLCγ2 activation. HeLa cells subjected to RAC1 KD or mock treatment were serum-starved for 16 h, stimulated with high-dose EGF (100 ng/ml), and harvested for immunoblot analysis at the indicated times. PLCγ1/2 activation and RAC1 KD efficiency were assessed. Vinculin, loading control. Lower, time course of PLCγ1/2 phosphorylation relative to total protein, normalized to stimulated control (2 min time point); n = 3. Student’s paired t-test: *P < 0.05, ns. **B.** RAC1 is required for EGF-induced PM-localized Ca^2+^ signaling. HeLa cells expressing the PM-GCaMP6f Ca^2+^ sensor were subjected to the indicated KDs and stimulated with high-dose EGF (100 ng/ml). Upper, representative Ca^2+^ traces (490/406 nm emission ratio). Lower, box plots of Ca^2+^ response AUC. N (cells): Control = 297, PLCγ2 KD = 271, RAC1 KD = 323; n = 2. One-way ANOVA: ****P < 0.0001. **C.** Direct interaction between PLCγ2 with RAC1 is required for EGFR-NCE. HeLa cells expressing HA-tagged siRNA-resistant PLCγ2-WT, the RAC1-binding-deficient PLCγ2 F897Q mutant, or empty vector (EV) were subjected to PLCγ2 KD or mock treatment and stimulated with Alexa Fluor-555 EGF (100 ng/ml) for 5 min. Upper, representative confocal images: CD147, green; EGF, red; DAPI, blue. Scale bar, 10 µm. Lower, quantification of internalized CD147, expressed as percentage of the EV control (box plots of mean integrated fluorescence intensity). N (fields): EV-PLCγ2 KD = 27, EV+PLCγ2 KD = 28, PLCγ2 WT-PLCγ2 KD = 28, PLCγ2 WT+PLCγ2 KD = 28, PLCγ2 F897Q-PLCγ2 KD = 28, PLCγ2 F897Q+PLCγ2 KD = 28; n = 3. One-way ANOVA: ****P < 0.0001; ns.

Structural studies have shown that RAC binds the split-PH domain of PLCγ2 through a hydrophobic cleft containing residues K862, V893 and F897, which are not conserved in PLCγ1 and are critical for RAC-dependent activation of PLCγ2^22^. The PLCγ2 F897Q substitution disrupts RAC binding and prevents RAC-mediated activation^22^. To determine whether this interaction is required for EGFR-NCE, we generated stable HeLa cell lines expressing siRNA-resistant PLCγ2-WT or the RAC-binding-defective PLCγ2-F897Q mutant at comparable levels (**Fig. S5C**) and assessed their ability to rescue CD147 internalization following PLCγ2 depletion. Whereas re-expression of PLCγ2-WT fully restored CD147 endocytosis, the PLCγ2-F897Q mutant failed to do so (**Fig. 5C**), supporting a requirement for the RAC-binding interface of PLCγ2 in EGFR-NCE.

### EGFR-NCE distinguishes cellular responses to EGF and TGFα

We next asked whether engagement of EGFR-NCE is a general feature of EGFR ligands or is ligand-selective. We focused on TGFα, a high-affinity EGFR ligand that binds the receptor with an affinity similar to that of EGF (K_d_ EGF=1.9nM vs. TGFα= 9.2nM)^23^, but elicits a stronger mitogenic response at equivalent concentrations^6, 24^.

Dose-response analysis during the initial phase of receptor activation (2 min) revealed that EGF induced slightly but consistently higher EGFR phosphorylation than TGFα, whereas phosphorylation of the major downstream signaling effectors was comparable between the two ligands (**Fig. 6A**). However, following prolonged stimulation with saturating ligand concentrations, TGFα produced more sustained activation of downstream signaling effectors, including MAPK and AKT, compared with EGF, accompanied by delayed EGFR degradation (**Fig. 6B**). These findings are in line with the stronger proliferative response of TGFα relative to EGF.

**Figure 6.**
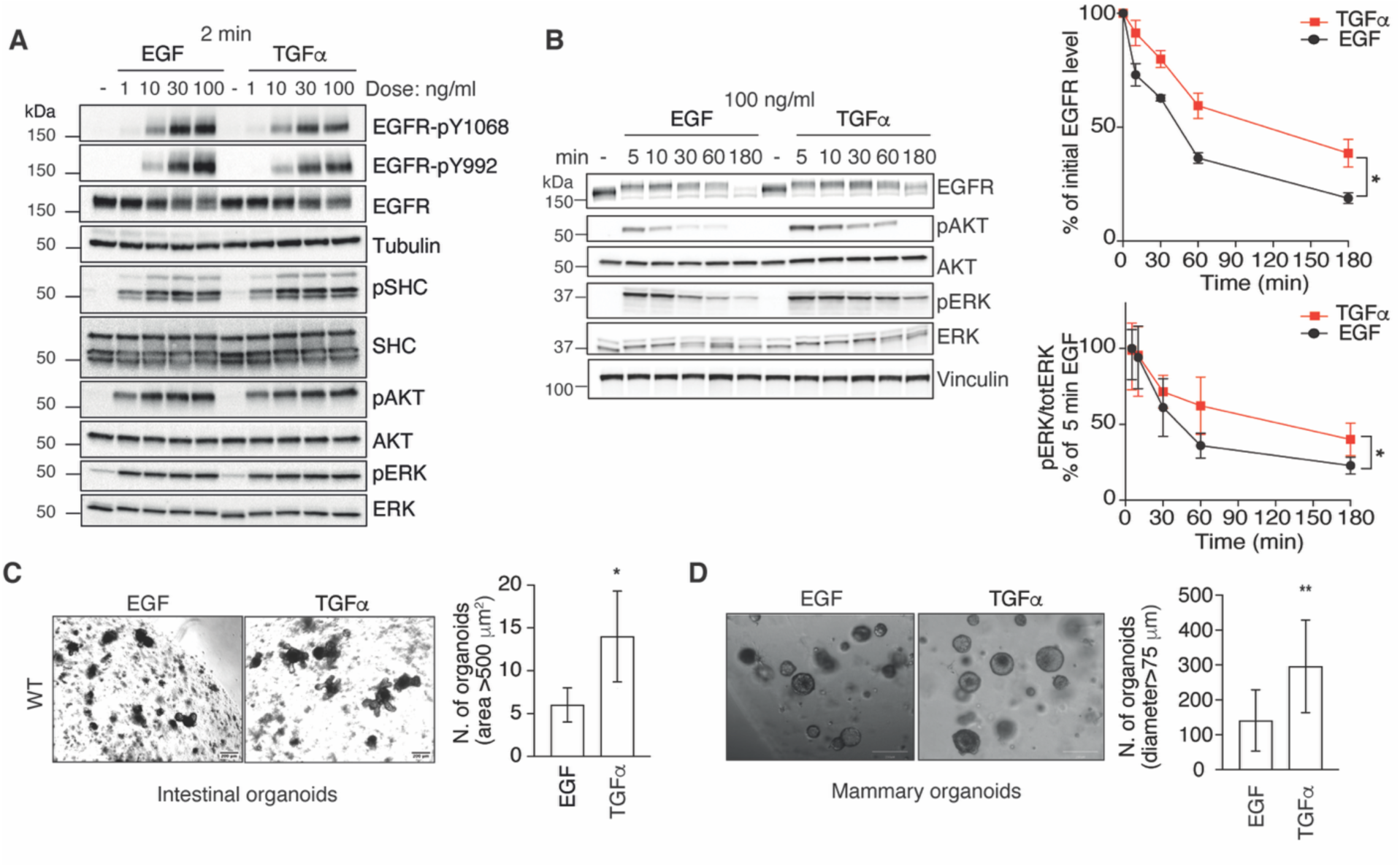
TGFα promotes sustained EGFR signaling and RTN3-independent organoid growth. **A.** EGF and TGFα induce comparable early EGFR pathway activation. HeLa cells were serum-starved for 16 h, stimulated with increasing concentrations of EGF or TGFα for 2 min, and harvested for immunoblot analysis of EGFR and downstream effector activation. Tubulin, loading control. MW markers shown on the left. **B.** Prolonged TGFα stimulation promotes sustained EGFR activation and reduced EGFR degradation relative to EGF. HeLa cells were serum-starved for 16 h, stimulated with high-dose EGF or TGFα (100 ng/ml), and harvested for immunoblot analysis at the indicated times. Vinculin, loading control. MW markers shown on the left. Right, upper, time course of total EGFR levels, expressed as percentage of the unstimulated control (mean ± SEM); n = 3. Lower, time course of ERK activation (pERK/total ERK ratio), expressed as percentage of the EGF-stimulated sample at 5 min (mean ± SEM); n = 3. AUC was calculated for the three independent experiments, and statistical significance was assessed by the Ratio paired t-test: *P < 0.05. **C.** TGFα promotes enhanced growth of intestinal organoids relative to EGF. Organoids derived from C57BL/6 mice were grown for 7 days in Matrigel in the presence of EGF or TGFα (50 ng/ml). Left, representative brightfield images of organoids at day 5. Right, quantification of the number of organoids with an area >500 µm^2^ (mean ± SD); N = 3. Student’s unpaired t-test: *P < 0.05. **D.** TGFα promotes enhanced growth of mammary organoids relative to EGF. Organoids derived from FVB mice were grown for 8 days in Matrigel-Collagen I mix in the presence of EGF or TGFα (50 ng/ml). Left, representative brightfield images of organoids at day 8. Scale bar, 200 μm. Right, quantification of the number of organoids with a diameter >75 µm (mean ± SD); N=9; n=3.

To determine whether these signaling differences translate into distinct biological outcomes, we compared the effects of EGF and TGFα on the growth of primary mouse intestinal organoids. Organoid growth requires the presence of EGF in the medium and is dependent on EGFR kinase activity, as pharmacological inhibition of the receptor abolished this response (**Fig. S6A**)^25^. Substituting EGF with TGFα increased the number of intestinal organoids above the prespecified area threshold (**Fig. 6C**). A similar increase in the number of mammary organoids above the diameter threshold was observed after TGFα exposure, indicating that this effect is not restricted to intestinal epithelium (**Fig. 6D and Fig. S6B**).

Strikingly, organoids derived from RTN3-KO mice, which are defective in EGFR-NCE, yielded more organoids above the prespecified size thresholds in response to both TGFα and EGF, resembling the phenotype of WT organoids stimulated with TGFα (**Fig. 6C,D**). These findings are consistent with a role for the RTN3-dependent EGFR-NCE signaling platform in limiting EGF-driven organoid expansion. In the absence of this platform, the EGF response is associated with delayed receptor degradation and sustained proliferative signaling, possibly as a result of endocytic recycling to the cell surface. Furthermore, they suggest that the greater organoid growth response to TGFα arises, at least in part, from its inability to engage this regulatory platform.

### TGFα selectively engages CME while failing to activate EGFR-NCE

We next investigated whether the greater mitogenic activity of TGFα reflects a reduced ability to activate the RAC1-PLCγ2-IP3R signaling axis and, consequently, to engage the EGFR-NCE organelle platform. Time-course analysis in HeLa cells revealed that stimulation with a high concentration of TGFα (100 ng/ml) induced substantially weaker PLCγ2 phosphorylation compared with EGF (**Fig. 7A**), whereas PLCγ1 phosphorylation was reduced to a much lesser extent. These effects were paralleled by reduced phosphorylation of the EGFR itself following TGFα stimulation relative to EGF (**Fig. 7A and Fig. S6C,D**), whereas activation of other canonical EGFR signaling effectors, including AKT, SHC and MAPK, was comparable between the two ligands throughout the time course.

**Figure 7.**
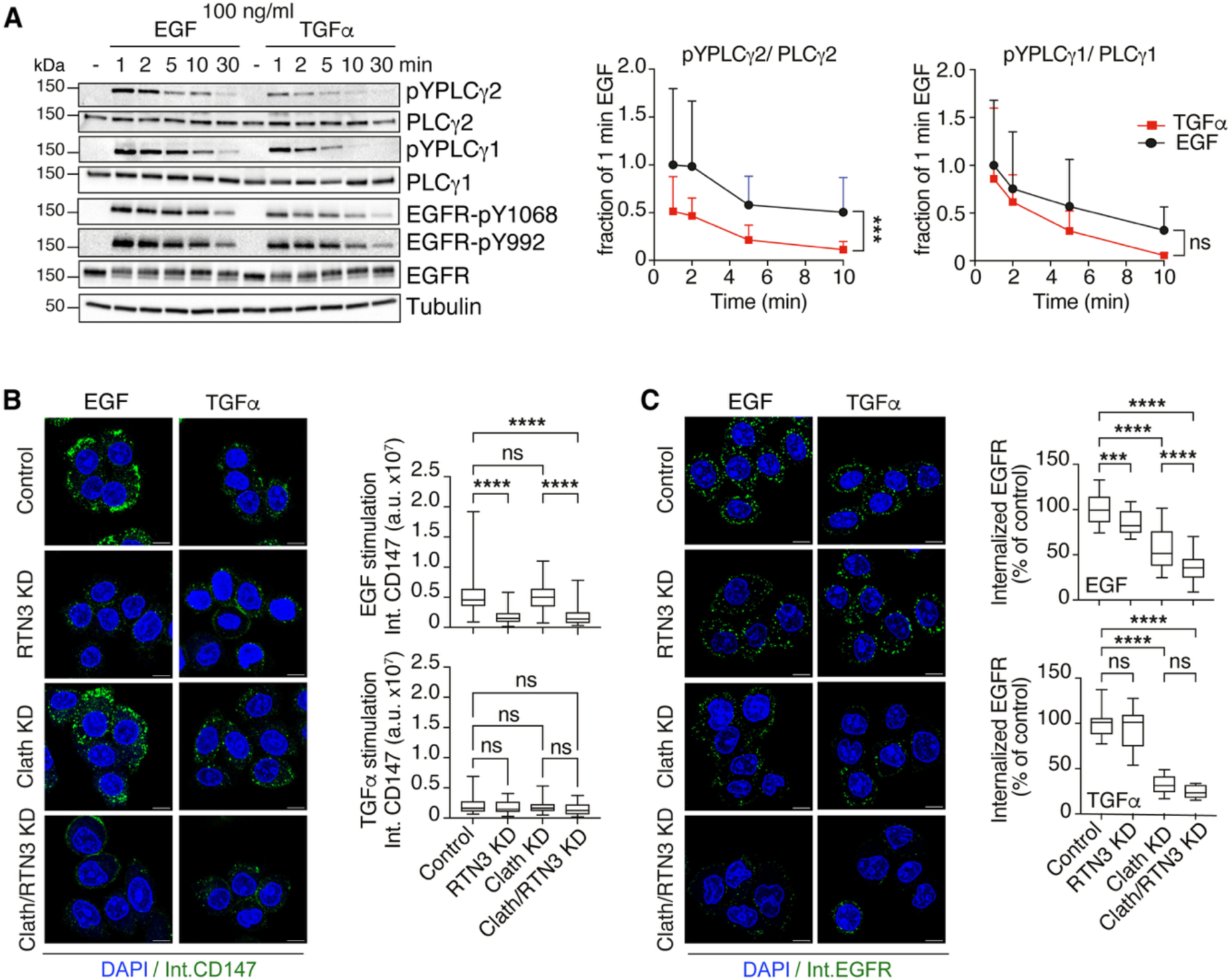
Differential PLCγ2 activation by TGFα and EGF correlates with selective engagement of CME versus EGFR-NCE. **A.** TGFα is a less potent activator of PLCγ2 than EGF. Left, HeLa cells were serum-starved for 16 h, stimulated with high-dose EGF or TGFα (100 ng/ml), and harvested for immunoblot analysis at the indicated times. Tubulin, loading control. MW markers shown on the left. Right, time course of PLCγ1/2 activation (pPLCγ1/2/total PLCγ1/2 ratio) normalized to the EGF-stimulated sample at 1 min (mean ± SEM); n = 3. AUC was calculated for the three independent experiments, and statistical significance was assessed by the Ratio paired t-test: ***P < 0.001; ns. **B.** TGFα does not activate RTN3-dependent EGFR-NCE. CD147 internalization was monitored by IF in HeLa cells subjected to the indicated KDs or mock treatment and stimulated with high-dose EGF (100 ng/ml) or TGFα (100 ng/ml) for 8 min. Left, representative confocal images: internalized CD147 (Int.CD147), green; DAPI, blue. Scale bar, 10 µm. Right, quantification of internalized CD147 in EGF-(upper) or TGFα-stimulated (lower) cells: box plot of mean integrated fluorescence intensity in arbitrary units (a.u.). EGF, N (fields), Control = 37, RTN3 KD = 40, Clath KD = 39, Clath/RTN3 KD = 37; n = 3. TGFα, N (fields), Control = 39, RTN3 KD = 39, Clath KD = 39, Clath/RTN3 KD = 40; n = 3. One-way ANOVA: ****P < 0.0001, ns. **C.** TGFα induces EGFR internalization via CME but not EGFR-NCE. EGFR internalization was monitored by IF in HeLa cells subjected to the indicated KDs or mock treatment and stimulated with high-dose of EGF or TGFα (100 ng/ml) for 8 min. Left, representative confocal images, internalized EGFR (Int.EGFR), green; DAPI, blue. Scale bar, 10 µm. Right, quantification of internalized EGFR in EGF-(upper) or TGFα-(lower) stimulated cells, expressed as percentage of control cells (box plot of mean integrated fluorescence intensity). EGF, N (fields), Control = 25, RTN3 KD = 26, Clath KD = 29, Clath/RTN3 KD = 29; n = 3. TGFα, N (fields), Control = 15, RTN3 KD=12, Clath KD = 15, Clath/RTN3 KD = 15; n = 2. One-way ANOVA: ****P < 0.0001, ***P < 0.001, ns.

The attenuated activation of PLCγ2 by TGFα was accompanied by a marked impairment in EGFR-NCE. Internalization of the NCE cargo CD147 was substantially reduced in TGFα-stimulated cells compared with EGF-stimulated cells, and, unlike EGF-induced uptake, was insensitive to RTN3 depletion (**Fig. 7B**). To directly examine the endocytic route of ligand-bound EGFR, we tracked receptor internalization using a live-cell labeling approach with an EGFR antibody that does not interfere with receptor activation (clone 13A9)^19^. As expected, EGF promoted EGFR internalization through both NCE and CME, as receptor uptake was partially inhibited by either RTN3 or clathrin KD and further reduced by combined depletion of both proteins (**Fig. 7C**). In contrast, TGFα-stimulated EGFR was internalized almost exclusively through CME, as clathrin KD nearly abolished receptor uptake, whereas RTN3 KD had no detectable effect (**Fig. 7C**). Thus, despite broadly comparable activation of several downstream EGFR effectors, TGFα fails to efficiently activate the PLCγ2-dependent NCE pathway and instead directs EGFR predominantly into the CME pathway.

### TGFα fails to activate the EGFR-NCE organelle platform and its associated cellular responses

Given that TGFα fails to engage EGFR-NCE and only weakly activates PLCγ2, we hypothesized that TGFα-bound EGFR would also be unable to promote the CTxB-positive PM regions associated with NCE. Consistent with this hypothesis, only EGF increased CTxB staining at the PM (**Fig. 8A**). This was accompanied by defective assembly of the EGFR-NCE organelle platform as assessed by EM (**Fig. 8B**). TGFα also elicited a distinct subplasmalemmal Ca²⁺ response compared with EGF, characterized by a delayed onset (∼2 min after stimulation *vs.* ∼30 sec for EGF) and a shorter-lived oscillatory response (**Fig. 8C and Fig. S6E**). These altered kinetics are consistent with the defective formation of PM-ER contact sites, which likely limits efficient coupling between PM-generated IP3 and ER-localized IP3 receptors.

**Figure 8.**
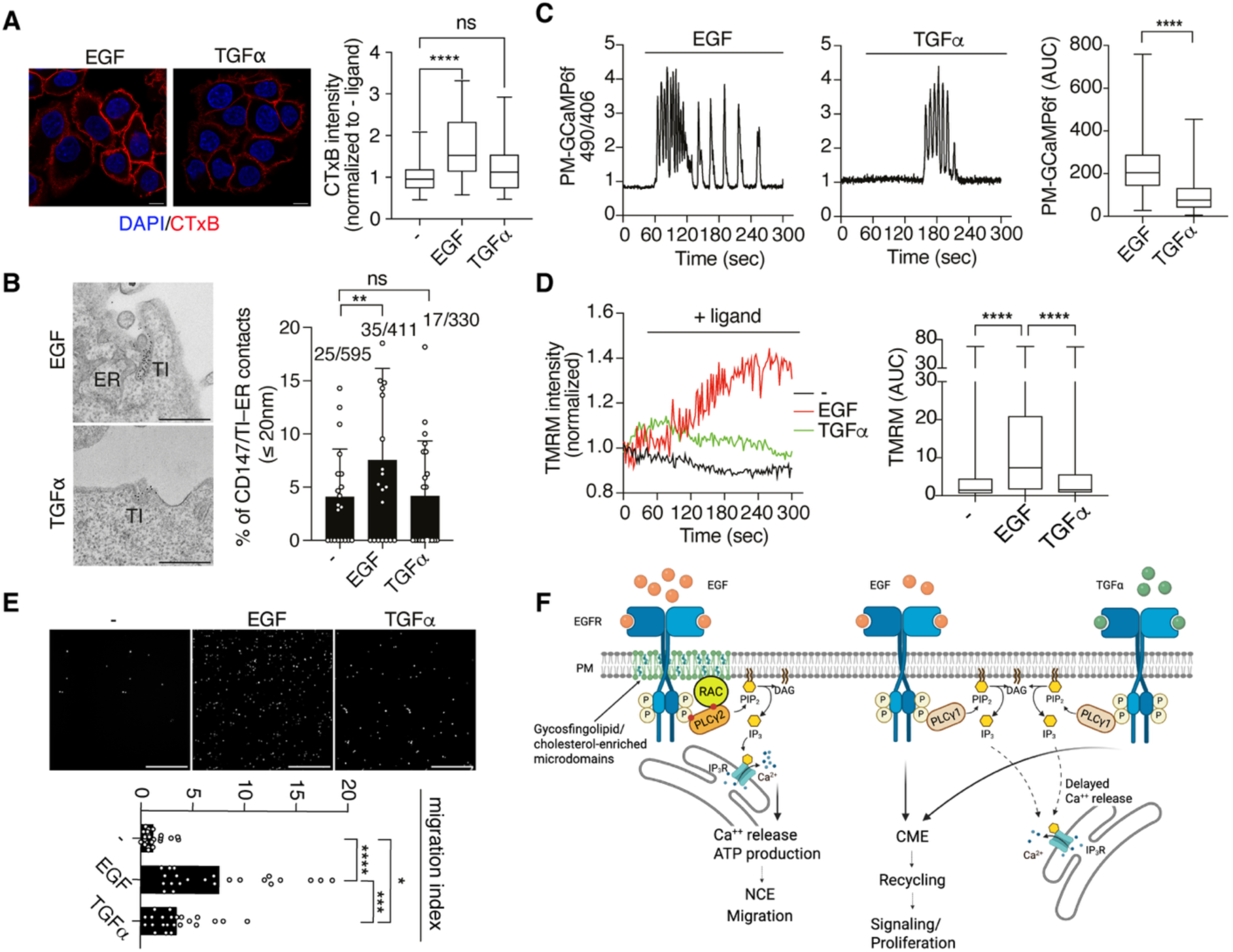
TGFα fails to activate the EGFR-NCE organelle platform and its associated Ca^2+^, mitochondrial, and migratory responses. **A.** TGFα does not induce the formation of GM1-rich PM microdomains. HeLa cells were serum-starved for 2 h and stimulated or not (−) with high-dose EGF or TGFα (100 ng/ml) for 1 min. Cells were fixed, incubated with Alexa Fluor 555-conjugated CTxB (285 ng/ml), permeabilized and analyzed by confocal microscopy. Left, representative IF images: CTxB, red; DAPI, blue. Scale bar, 10 µm. Right, quantification of CTxB binding as signal intensity normalized to unstimulated control (box plot of mean integrated fluorescence intensity). N (fields): No ligand = 30, EGF = 28, TGFα=28 (n = 3). One-way ANOVA: **P < 0.01; ****P < 0.0001; ns. **B.** TGFα does not promote ER recruitment of CD147-positive endocytic structures. ER proximity of CD147-positive endocytic structures was assessed by EM morphometry using gold-labeled CD147 in HeLa cells stimulated or not (−) with high-dose EGF or TGFα (30 ng/ml) for 5 min. Left, representative EM images. Scale bar, 500 nm. Right, percentage of CD147-positive structures in contact with the ER (≤ 20 nm) shown as mean ± SD. N (cells): No ligand=20, EGF =20, TGFα=21. One-way ANOVA, **P < 0.01; ns. **C.** TGFα elicits delayed and more transient PM-localized Ca^2+^ response compared to EGF. HeLa cells expressing the PM-GCaMP6f Ca^2+^ sensor were stimulated with high-dose EGF or TGFα (100 ng/ml). Left, representative Ca^2+^ traces (490/406 nm emission ratio). Right, box plots of Ca^2+^ response AUC. N (cells): EGF = 157, TGFα = 139; n =3. Unpaired t-test two-tailed; ****P < 0.0001. **D.** TGFα fails to efficiently stimulate mitochondrial bioenergetics. Mitochondrial membrane potential was monitored by TMRM fluorescence in HeLa cells stimulated or not (−) with high-dose EGF or TGFα (100 ng/ml). Left, representative time course of TMRM fluorescence. Right, box plots of AUC. N (cells): No ligand = 564, EGF = 802, TGFα = 731; n = 3. Unpaired t-test two-tailed: ****P < 0.0001. **E.** TGFα is a weak inducer of cell migration relative to EGF. HeLa cells were serum-starved for 16 h and assessed for migratory using a Transwell assay over 24 h. Cells were seeded in the upper chamber, while high-dose EGF or TGFα (100 ng/ml), or serum-free medium, was added to the bottom chamber. Lower, quantification of migrating cells per field of view, normalized to unstimulated control (mean ± SD); n = 3. One-way ANOVA: ****P < 0.0001; ***P < 0.001; *P < 0.05. **F.** Schematic illustrating the differential engagement of CME and EGFR-NCE by low- and high-dose EGF and by TGFα, driven by differential activation of the RAC1–PLCγ2–IP3R signaling axis.

The attenuated Ca²⁺ response induced by TGFα did not produce a significant increase in TMRM fluorescence (Fig. 8D) (**Fig. 8D**). Since local mitochondrial ATP production at PM-ER-mitochondria contact sites is required for cortical actin remodeling and EGF-induced cell migration^5^, we next assessed cell migration using Transwell assays. Consistent with its inability to assemble the EGFR-NCE platform, TGFα was significantly less effective than EGF at promoting cell migration (**Fig. 8E**).

Collectively, these findings support a model in which differential engagement of the RAC1-PLCγ2-IP3R signaling axis by distinct EGFR ligands determines whether the receptor engages the NCE organelle platform. Efficient engagement of this pathway by EGF couples receptor signaling to localized Ca²⁺ signaling, a mitochondrial membrane-potential response, and migratory behavior, whereas its inefficient engagement by TGFα favors internalization via CME, delayed receptor degradation and sustained proliferative signaling.

## Discussion

The ability of a single receptor to generate distinct cellular responses is attributed to differences in ligand type and concentration, receptor conformation and spatial organization, signaling dynamics and intracellular trafficking. Here, we connect these levels of regulation by implicating a RAC1-PLCγ2 signaling axis in the engagement of the EGFR-NCE organelle platform and in ligand-dependent biological outputs. Our data support a model in which high-dose EGF, but not TGFα, promotes the assembly of specialized PM domains associated with RTN3-dependent ER-PM contact sites. Within these domains, RAC1 is required for efficient PLCγ2 phosphorylation and for IP3R-dependent Ca²⁺ release, mitochondrial Ca²⁺ uptake, and the associated change in mitochondrial membrane potential. This signaling cascade is required for efficient NCE tubular fission and is associated with receptor downregulation and enhanced cell migration. Conversely, the weaker engagement of this pathway by TGFα is associated with predominantly CME-dependent receptor internalization, delayed receptor degradation, and sustained mitogenic signaling (**Fig. 8F**). Thus, our findings support a model in which differential assembly of an endocytic organelle platform contributes to EGFR ligand bias.

A major finding of this study is the selective role of PLCγ2 over PLCγ1 in regulating EGFR-NCE. PLCγ1 is generally considered the predominant PLCγ enzyme acting downstream of receptor tyrosine kinases in epithelial cells, whereas PLCγ2 has been studied mainly in hematopoietic and immune cells. Surprisingly, both family members become phosphorylated after high-dose EGF stimulation, yet only PLCγ2 is required for subplasmalemmal Ca²⁺ signaling, mitochondrial activation, and NCE. Our data suggest that this functional specificity is associated with distinct spatial organization rather than simply with receptor-induced phosphorylation. PLCγ2 shows greater association than PLCγ1 with PM regions juxtaposed to RTN3-positive cortical ER, potentially positioning PIP2 hydrolysis close to IP3R (Fig. 3C). These findings are consistent with the emerging concept that spatial compartmentalization of signaling enzymes can contribute to signaling specificity.

Our data further support RAC1 as an important determinant of PLCγ2 function in this pathway. Structural and biochemical studies established that activated RAC binds the split PH domain of PLCγ2, but not PLCγ1, through a hydrophobic interface involving residues including Phe897, thereby stimulating PLCγ2 catalytic activity^13, 22^. Consistent with these observations, the RAC-binding-defective mutant PLCγ2-F897Q failed to rescue NCE in PLCγ2-depleted cells, supporting a requirement for RAC-dependent PLCγ2 function in epithelial cells. Beyond directly activating PLCγ2, RAC1 appears to organize the membrane environment in which EGFR-NCE is initiated. RAC1 depletion reduces PM-associated CTxB signal, impairs PLCγ2 phosphorylation, and prevents formation of RTN3-dependent ER-PM contacts. Together, these findings support a model in which EGF-dependent RAC1 function promotes membrane organization and ER-PM contact formation, while engagement of PLCγ2 by EGFR and RAC1 may facilitate its selective activation despite its relatively low abundance. Thus, a signaling module originally characterized in immune cells may also operate in epithelial cells to organize a ligand-regulated endocytic platform.

The RAC1-PLCγ2-IP3R pathway propagates beyond the ER and affects mitochondrial Ca²⁺ handling and membrane potential. Inhibition of this pathway suppresses mitochondrial Ca²⁺ signals and the rapid increase in mitochondrial membrane potential induced by EGF, supporting coupling between EGFR signaling and a mitochondrial bioenergetic response^5^. These observations may also help reconcile the apparently opposing functions of this platform. On the one hand, EGFR-NCE attenuates signaling by directing receptors toward lysosomal degradation, whereas CME sustains receptor recycling and prolonged signaling^3, 4^. On the other hand, previous work indicates that the same platform transiently promotes local mitochondrial ATP production to support actin remodeling and cell migration^5^. Rather than representing contradictory outputs, these processes coordinate a high-intensity growth factor response with its timely termination. PLCγ2 may therefore occupy a central position in this circuitry, linking receptor internalization to the Ca²⁺-dependent mitochondrial response associated with cell migration.

An important implication of our findings is that differential engagement of the PLCγ2-dependent platform contributes to EGFR ligand bias. EGF and TGFα bind EGFR with similar affinity and induce comparable early receptor phosphorylation, yet they produce distinct trafficking and biological responses. EGF promotes receptor ubiquitination and lysosomal degradation, whereas TGFα preferentially supports receptor recycling and sustained proliferation^26, 27^. These differences have been attributed to the greater acid sensitivity of the TGFα-EGFR complex, which favors ligand dissociation within endosomes and receptor recycling^7–9, 28^, and to the recruitment of specific adaptors at the late endosomes^10^. Our results do not contradict this model but suggest that ligand-dependent differences may also emerge at the PM. TGFα is associated with weaker PLCγ2 activation, reduced assembly of the organelle platform, and attenuated Ca²⁺ and mitochondrial membrane-potential responses. Thus, EGF and TGFα begin to engage distinct signaling and trafficking programs before endosomal sorting.

How ligand identity is translated into selective RAC1 and PLCγ2 activation remains an open question. However, our data are in line with structural and biophysical studies showing that different EGFR ligands stabilize distinct receptor conformations and juxtamembrane organizations, influencing receptor clustering, lateral mobility to PM microdomains, and downstream signaling^8, 9, 29–31^. It is therefore plausible that EGF-bound EGFR more efficiently adopts an organization that favors RAC1-dependent membrane remodeling and PLCγ2 engagement. Consistent with this idea, RAC1 depletion reduces PM-associated CTxB signal and ER-PM contact formation, providing a potential link between ligand-dependent receptor organization and assembly of the NCE machinery. Whether RAC1 directly senses specific EGFR conformations or is activated through ligand-specific GEFs remains to be established. Finally, our biological data suggest that differential endocytic routing may contribute to the higher mitogenic activity of TGFα compared with EGF. TGFα increases the number of mammary and intestinal organoids above the predefined size thresholds compared with EGF, whereas EGF-treated RTN3-deficient organoids show a similar trend. These observations are consistent with a role for the NCE platform in restraining the mitogenic output of EGF. Conversely, weaker activation of the PLCγ2-dependent pathway by TGFα is associated with altered Ca²⁺ dynamics, lack of a detectable increase in mitochondrial membrane potential, and reduced migration in the Transwell assay.

In conclusion, our study supports PLCγ2 as a receptor-proximal effector linking RAC1-dependent membrane organization to IP3R-mediated Ca²⁺ signaling at the EGFR-NCE organelle platform. More broadly, our findings suggest that endocytic organelle platforms can contribute actively to the decoding of ligand bias rather than acting solely as passive consequences of receptor activation. By integrating receptor organization, intracellular trafficking, and organelle communication, the RAC1-PLCγ2 axis may help translate ligand identity into distinct biological outputs, providing a framework for understanding how closely related EGFR ligands generate different cellular responses.

## Acknowledgements

We thank Rosalind Gunby for critically editing the manuscript. We thank Alberto Gobbi and the mouse facility (Cogentech Società Benefit Srl, Milan) for animal husbandry. We thank all IEO Technological Units. We thank Simona Rodighiero and the Imaging Unit, and Simona Ronzoni and the Flow Cytometry Unit for their technical support. We thank Marika Zanotti, Diego Pasini, Michela Bruzzi, and Marina Mapelli for sharing organoid mouse culture protocols and for their valuable advice and technical guidance. We thank the Department of Oncology and Hematology-Oncology at the University of Milan for the administrative support along the project.

## Author contributions

Investigation: GJ, DM, SF, GM, CT, AFB, MQ, MC, AR, GC, EB, SP, DC, FB. Methodology: SF, AR, SP, MF, MB. Data analysis: GJ, SF, DC, FB, MB, SP. Visualization: GJ, SS, CT, MGM. Resources: SS, MB, PPDF. Funding acquisition: SS, PPDF. Supervision: SS, PPDF, MF, PP, MB. Conceptualization: SS, PPDF. Writing – original draft: SS; Writing – review & editing: SS, PPDF, GJ, CT, MGM.

## Data and materials availability

All data are available in the main text or the supplementary materials. Reagents and materials used in this study are either commercially available or can be obtained upon request.

## References

1. Singh, B., Carpenter, G. & Coffey, R.J. EGF receptor ligands: recent advances. F1000Res 5 (2016).

2. Jendrisek, G., Mesa, D., Conte, A., Malabarba, M.G. & Sigismund, S. Beyond the membrane: rethinking EGFR signaling in physiology and cancer. Cell Mol Life Sci 83, 109 (2026).

3. Sigismund, S. et al. Clathrin-mediated internalization is essential for sustained EGFR signaling but dispensable for degradation. Dev Cell 15, 209–219 (2008).

4. Caldieri, G. et al. Reticulon 3-dependent ER-PM contact sites control EGFR nonclathrin endocytosis. Science 356, 617–624 (2017).

5. Mesa, D. et al. A tripartite organelle platform links growth factor receptor signaling to mitochondrial metabolism. Nat Commun 15, 5119 (2024).

6. Yarden, Y. & Sliwkowski, M.X. Untangling the ErbB signalling network. Nat Rev Mol Cell Biol 2, 127–137 (2001).

7. Ebner, R. & Derynck, R. Epidermal growth factor and transforming growth factor-alpha: differential intracellular routing and processing of ligand-receptor complexes. Cell Regul 2, 599–612 (1991).

8. Doerner, A., Scheck, R. & Schepartz, A. Growth Factor Identity Is Encoded by Discrete Coiled-Coil Rotamers in the EGFR Juxtamembrane Region. Chem Biol 22, 776–784 (2015).

9. Sinclair, J.K.L., Walker, A.S., Doerner, A.E. & Schepartz, A. Mechanism of Allosteric Coupling into and through the Plasma Membrane by EGFR. Cell Chem Biol 25, 857–870 e857 (2018).

10. Francavilla, C. et al. Multilayered proteomics reveals molecular switches dictating ligand-dependent EGFR trafficking. Nat Struct Mol Biol 23, 608–618 (2016).

11. Sigismund, S. et al. Threshold-controlled ubiquitination of the EGFR directs receptor fate. EMBO J 32, 2140–2157 (2013).

12. Wilson, B.S., Pfeiffer, J.R., Surviladze, Z., Gaudet, E.A. & Oliver, J.M. High resolution mapping of mast cell membranes reveals primary and secondary domains of Fc(epsilon)RI and LAT. J Cell Biol 154, 645–658 (2001).

13. Walliser, C. et al. rac regulates its effector phospholipase Cgamma2 through interaction with a split pleckstrin homology domain. J Biol Chem 283, 30351–30362 (2008).

14. Walliser, C. et al. Rac-mediated Stimulation of Phospholipase Cgamma2 Amplifies B Cell Receptor-induced Calcium Signaling. J Biol Chem 290, 17056–17072 (2015).

15. Innocenti, M. et al. Phosphoinositide 3-kinase activates Rac by entering in a complex with Eps8, Abi1, and Sos-1. J Cell Biol 160, 17–23 (2003).

16. Zhu, G. et al. An EGFR/PI3K/AKT axis promotes accumulation of the Rac1-GEF Tiam1 that is critical in EGFR-driven tumorigenesis. Oncogene 34, 5971–5982 (2015).

17. del Pozo, M.A. et al. Integrins regulate Rac targeting by internalization of membrane domains. Science 303, 839–842 (2004).

18. Ridley, A.J., Paterson, H.F., Johnston, C.L., Diekmann, D. & Hall, A. The small GTP-binding protein rac regulates growth factor-induced membrane ruffling. Cell 70, 401–410 (1992).

19. Sigismund, S. et al. Clathrin-independent endocytosis of ubiquitinated cargos. Proc Natl Acad Sci U S A 102, 2760–2765 (2005).

20. Simons, K. & Gerl, M.J. Revitalizing membrane rafts: new tools and insights. Nat Rev Mol Cell Biol 11, 688–699 (2010).

21. Chang, C.L. et al. Feedback regulation of receptor-induced Ca2+ signaling mediated by E-Syt1 and Nir2 at endoplasmic reticulum-plasma membrane junctions. Cell Rep 5, 813–825 (2013).

22. Bunney, T.D. et al. Structural insights into formation of an active signaling complex between Rac and phospholipase C gamma 2. Mol Cell 34, 223–233 (2009).

23. Jones, J.T., Akita, R.W. & Sliwkowski, M.X. Binding specificities and affinities of egf domains for ErbB receptors. FEBS Lett 447, 227–231 (1999).

24. Luetteke, N.C. & Lee, D.C. Transforming growth factor alpha: expression, regulation and biological action of its integral membrane precursor. Semin Cancer Biol 1, 265–275 (1990).

25. Sato, T. et al. Single Lgr5 stem cells build crypt-villus structures in vitro without a mesenchymal niche. Nature 459, 262–265 (2009).

26. Waterman, H., Sabanai, I., Geiger, B. & Yarden, Y. Alternative intracellular routing of ErbB receptors may determine signaling potency. J Biol Chem 273, 13819–13827 (1998).

27. Roepstorff, K. et al. Differential effects of EGFR ligands on endocytic sorting of the receptor. Traffic 10, 1115–1127 (2009).

28. Decker, S.J. Epidermal growth factor and transforming growth factor-alpha induce differential processing of the epidermal growth factor receptor. Biochem Biophys Res Commun 166, 615–621 (1990).

29. Needham, S.R. et al. EGFR oligomerization organizes kinase-active dimers into competent signalling platforms. Nat Commun 7, 13307 (2016).

30. Ronan, T. et al. Different Epidermal Growth Factor Receptor (EGFR) Agonists Produce Unique Signatures for the Recruitment of Downstream Signaling Proteins. J Biol Chem 291, 5528–5540 (2016).

31. Freed, D.M. et al. EGFR Ligands Differentially Stabilize Receptor Dimers to Specify Signaling Kinetics. Cell 171, 683–695 e618 (2017).

