## Supplementary Material for "Decoding EGFR ligand bias through an endocytic organelle platform"

#### Supplementary Materials

#### Materials and Methods

##### Reagents

Throughout the manuscript, EGF (Thermo Fisher Scientific, AF-100-15) and TGF-α (Thermo Fisher Scientific, 100-16A) were used for receptor stimulation at the indicated concentrations. Alexa-labelled EGF (Thermo Fisher Scientific; Alexa Fluor™ 555, E35350; Alexa Fluor™ 647, E35351; and Alexa Fluor™ 488, E13345) was used at concentrations of 400 ng/mL or 1 µg/mL of the conjugated species, corresponding to actual EGF concentrations of approximately 40 ng/mL and 100 ng/mL, respectively. Alexa-Tf (Thermo Fisher Scientific, Alexa Fluor™ 488 Conjugate, T13342) was used at a concentration of approximately 50 µg/mL. Protein-A Gold 10 nm was obtained from Utrecht University; EM grade glutaraldehyde and paraformaldehyde were obtained from Electron Microscopy Sciences. Ruthenium red and secondary rabbit anti-mouse were from Sigma. Fluorescent secondary antibodies were obtained from Thermo Fisher Scientific (5 µg/mL). Gefitinib was from Merck Life Technology S.R.L (SML1657-10MG). Cholera Toxin Subunit B (Recombinant), Alexa Fluor™ 555 Conjugate (Thermo Fisher Scientific, C34776) was used at 285ng/ml.

| **Antibody** | **Producer** | **Clone or Epitope** | **Catalog number** | **Dilution** |
|---|---|---|---|---|
| AKT | Cell Signaling | | #9272 | 1:1000 |
| CD147 | BD | HIM 6 | 555961 | 1:300 |
| Clathrin | BD | clone 23 | 610499 | 1:1000 |
| EGFR | Genentech | Clone 13A9 | 13A9 | 1:20 000 (in vivo) |
| EGFR | In-house | 806 serum | | IF 1:200<br>WB 1:10000 |
| ERK1/2 | Sigma | ERK-1, 351-368 | M7927 | 1:5000 |
| GAPDH | Santa Cruz | 6C5 | sc-32233 | 1:3000 |
| HA | BioLegend | 16B12 | 901503 | WB, IF 1:5000 |
| IP3R | BD | Clone 2 | 610312 | 1:500 |
| pAKT | Cell Signaling | Thr308 | # 9275 | 1:500 |
| pERK1/2 | Cell Signaling | Thr202/Tyr204 | #9106 | 1:1000 |
| PLCγ1 | BD | Clone 10 | 558575 | 1:500 |
| PLCγ2 | R&D | 346404 | MAB3716 | 1:500 |
| pPLCγ1 | Cell Signaling | Tyr1783 | #2821 | 1:1000 |
| pPLCγ2 | R&D | Tyr759,<br>Clone #744757 | MAB7377 | 1:500 |
| Rac1 | Sigma | Clone 23A8 | 05-389 | 1:50 for IF<br>1:500 for WB |
| pSHC | Cell Signaling | Tyr239/240 | #2434 | 1:500 |
| pTyr | Millipore | 4G10 | 05-321MG | 1:1000 |
| pY 1045 EGFR | Cell Signaling | Tyr 1045 | #2237 | 1:1000 |
| pY 1068 EGFR | Cell Signaling | Tyr 1068, D7A5 | #3777 (XP) | WB 1:1000<br>IF 1:200 |
| pY 1173 EGFR | Cell Signaling | Tyr 1173, 53A5 | #4407 | 1:1000 |

| | | | | |
|---|---|---|---|---|
| pY 992 EGFR | Cell Signaling | Tyr 992 | #2235 | 1:1000 |
| RTN3 | In-house | Amino acids 1-47 | | 1:250 |
| SHC | BD | Clone 20 | #610878 | 1:500 |
| Tubulin | In-house | serum | | 1:500 |
| Vinculin | Sigma | clone hVIN-1 | V9131 | 1:5000 |

*WB: western blot; IF: immunofluorescence.*

**Constructs**

pSLIK-neo-SNAP25-GCaMP6f vector was cloned as described [1]. HeLa cells infected with the pSLIK-neo-SNAP25-GCaMP6f vector and selected in a medium containing 800 μg/mL neomycin for 7 days and SNAP25-GCaMP6f protein expression was induced one day before experiment at 500 ng/mL of Doxycycline.

pLVX-NEO-CMV-2mt-GCaMP6s and SNAP25-Aequorin were kindly provided by Dr. Massimo Bonora and Dr. Paolo Pinton from the University of Ferrara[2, 3].

Lentiviral expression plasmids encoding human PLCG1 (NM_002660.2) and PLCG2 (NM_002661.5) were custom designed and synthesized by VectorBuilder (Chicago, IL, USA). The constructs included a CMV promoter and a C-terminal HA tag. The following vectors were used: pLVX-CMV-hPLCG1-HA (VectorBuilder ID: VB200406-2275msy), pLVX-CMV-hPLCG2-HA (VectorBuilder ID: VB200406-2324qhm), pLVX-Puro-CMV-hPLCG2 (aa 1228–1234 mutation, resistant to PLCG2 oligo 5'-GGGAUGCCCUGGUUAAAGA-3')-HA (VectorBuilder ID: VB221117-1633bxc), and pLVX-Puro-CMV-hPLCG2(F897Q; aa 1228–1234 mutation, resistant to PLCG2 oligo 5'-GGGAUGCCCUGGUUAAAGA-3')-HA (VectorBuilder ID: VB221117-1636rxd). All plasmids were sequence-verified before use. HeLa cells and inducible clathrin-KD cells were then infected with the following lentiviral pLVX vectors: empty vector (EV), PLCγ1-HA, or PLCγ2-HA. GFP-MAPPER was a gift from Jen Liou (Addgene plasmid # 117721; http://n2t.net/addgene:117721;RRID:Addgene_117721)[4]. pSlik-GFP-MAPPER was synthetized by cloning the sequence of GFP-MAPPER insert inside a pEN_Tmcs vector (gift from Iain Fraser Addgene plasmid #25751; http://n2t.net/addgene:25751; RRID:Addgene_25751)[5] and subsequently recombining the insert in pSlik-Neo vector (gift from Iain Fraser, Addgene plasmid #25735; http://n2t.net/addgene:25735; RRID:Addgene_25735)[5]. HeLa cells were stably infected and selected with 800μg/ml neomycin and protein expression induced by 1μg/ml doxycycline overnight.

**Cell culture and transfection**

Human epithelial cervical cancer (HeLa) cells were cultured in Minimum Essential Medium (MEM, Sigma) supplemented with 10% FBS, 2 mM glutamine, sodium pyruvate 1 mM (Euroclone), 0.1 mM non-essential amino acids (Euroclone). The inducible clathrin KD HeLa clone 53 was grown in the same medium as HeLa cells but supplemented with 10% TET-System Approved FBS (PAA) instead of standard FBS. For ligand stimulation experiments, the indicated concentrations of human recombinant ligands were added to serum-starved medium for the indicated time points (details in the Reagents section). The pSLIK-SNAP25-GCaMP6f and pLVX-mitoGCaMP6s HeLa clones were grown in the same medium as HeLa cells supplemented with 750 μg/mL G418 (Adipogen Life Sciences). HaCaT (human immortalized keratinocytes) cells were cultured in DMEM medium (Thermo Fisher Scientific) supplemented with 10% FBS and 2 mM glutamine. All cells were cultured at 37ºC and 5% $CO_2$. Cells were passaged every 2-3 days to maintain sub-confluency. HeLa cells at 50–60% confluence were transfected using Lipofectamine 3000 (Thermo Fisher Scientific) according to the manufacturer's instructions. Experiments were performed 24-48 h after transfection.

All human cell lines were authenticated at each batch freezing by STR profiling (StemElite ID System, Promega). All cell lines were tested for mycoplasma at each batch freezing by PCR[6] and biochemical assay (MycoAlert, Lonza).

**RNA interference (RNAi)**

RNAi was performed with Lipofectamine RNAimax from Invitrogen, according to the manufacturer's protocol: for RTN3, Clathrin heavy chain, Rac1 and PLCγ1 KD, cells were transfected with 2 cycles of 8 nM of oligos; for PLCγ2 KD, cells were transfected with 2 cycles of 16 nM of oligos; for IP3-R1, IP3-R2, IP3-R3 KDs, 1 cycle of 8 nM of oligos. Mock-treated cells (incubated with Lipofectamine RNAimax in optimum medium) were used as control. Briefly, 1.5 x $10^6$ cells in a 10 cm culture plate were transfected in reverse: liposome complexes carrying siRNAs were formed for 15 min at room temperature (RT) and then added to trypsinized cells in suspension. The next day, cells were transfected in forward, adding the liposome/siRNA complexes to cells in adhesion. Experiments were performed 72 h after the second transfection. Oligonucleotide brands and sequences are listed below:

**List of RNAi oligo sequences**

| ID | Sequence |
|---|---|
| PLCγ1 Smart pool Dharmacon | 5'-GCAGCAAGAUCUACUACUC-3'<br>5'-GAUGGGAUGCCAGUUAUUU-3'<br>5'-GAGACAACCGCCUCUAGUU-3'<br>5'-GAAUGGAAUUUCGCCUGAA-3' |
| PLCγ1 Invitrogen | Catalog number 4392420 Assay ID s10631 |
| PLCγ1 siPOOL Diatech Lab Line srl | SI-G020-5335 |
| PLCγ2 Smart pool Dharmacon | Cat. #: J-008339-07<br>5'-GGGAUGCCCUGGUUAAAGA-3' oligo #2<br>Cat. #: J-008339-05 oligo #1<br>Cat. #: J-008339-08 oligo #3<br>Cat. #: J-008339-17 oligo #4 |
| RTN3 Stealth | 5'-CCCUGAAACUCAUUAUUCGUCUCUU-3' |
| Clathrin heavy chain Riboxx | 5'-UAAAUUUCCGGGCAAAGAGCCCCC-3' |
| Rac1 oligo Stealth | 5'-CCGGUGAAUCUGGGCUUAUGGGAUA-3' |
| Rac1 Smart Pool Dharmacon | 5'-GUGAUUUCAUAGCGAGUUU-3'<br>5'-GUAGUUCUCAGAUGCGUAA-3'<br>5'-AUGAAAGUGUCACGGGUAA-3'<br>5'-GAACUGCUAUUUCCUCUAA-3' |
| IP3-R1 Smart Pool Riboxx | 5'-UUAAACGAAAUGCUGCUCCCCC-3'<br>5'-AUAUGUAGAUGUUGUGCCCCC-3'<br>5'-AUAACUAGAUUGGAAGCCCCC-3' |
| IP3-R2 Smart Pool Riboxx | 5'-UUAUUUCUUUCUGAGCAGCCCCC-3'<br>5'-AUUGAUACAAGAAACGGCCCCC-3'<br>5'-AUCUUUAACAUAACAGGCCCCC-3' |
| IP3-R3 Smart Pool Riboxx | 5'-AUUAAGGUAAACUGAGUCCCCC-3'<br>5'-UUAUUCUUGUCAGUCCACGCCCCC-3'<br>5'-UAUAGAUGUUAUGGCCCACCCCC-3' |

**Immunoblot analysis**

Cells were lysed by adding RIPA buffer (50 mM Tris–HCl, 150 mM NaCl, 1 mM EDTA, 1% Triton X-100, 1% sodium deoxycholate, 0.1% SDS), plus protease inhibitor cocktail

(CALBIOCHEM) and phosphatase inhibitors (20 mM sodium pyrophosphate pH 7.5, 50 mM NaF, 2 mM PMSF, 10 mM Na3VO4 pH 7.5), directly to cell plates. Lysate was clarified by centrifugation at 13,200 rpm for 20 min at 4°C. Protein concentration was measured by the Bradford Assay (Biorad) and 30µg of protein was prepared in Laemmli 1x buffer (2% SDS, 62.5 mM Tris HCl, pH 6.8, 10% Glycerol, 0.1% Bromophenol blue, 5% β-Mercaptoethanol), with the exception of pPLCγ2 and total PLCγ2 (45µg of lysates). Protein samples were run on pre-cast gels with a gradient 4-20% (Biorad) and transferred to a nitrocellulose membrane (Biorad) using Trans-Blot (Biorad) according to manufacturer's instructions. Membranes were blocked with 5% milk diluted in tris buffered saline 0.1% Tween (TBS-T) and then incubated overnight with primary antibody according to the datasheet. Following 3 washes with TBS-T, membranes were incubated with the appropriate secondary antibody conjugated to horseradish peroxidase. After 3 more washes, the signal was detected at Chemidoc (Biorad) using ECL from Amersham, Biorad or Femto.
Western blot band intensities were quantified using Fiji (ImageJ, National Institutes of Health, Bethesda, MD, USA). Band intensity was measured using the Gel Analyzer tool. Individual lanes were selected, and intensity profiles were generated using the Plot Lanes function. The integrated density of each band was determined by measuring the area under the corresponding peak after background subtraction. Protein expression was normalized to the corresponding loading control and phospho-to-total protein ratio, as appropriate. Relative protein levels were expressed as fold change relative to the control sample.

**CD147/EGF/Tf internalization assays**
For antibody internalization assays, cells plated on glass coverslips were first incubated with anti-CD147 or anti-EGFR (13A9) antibody for 30 min at 4˚C. Cells were then stimulated with high doses of different ligands (Alexa-EGF, EGF, TGFα) at 37°C for the indicated times. After internalization, cells were acid wash-treated (100 mM Glycine-HCl) pH 2.2 on ice for 3 times, 45 sec/each, fixed in 4% paraformaldehyde and processed for IF. After permeabilization, proteins of interest were labeled with adequate primary antibodies in 1% BSA/PBS, followed by incubation with specific secondary antibodies. Alexa-488 secondary antibody was used to label the internalized CD147 or EGFR antibody.
Images were obtained using a Leica TCS SP8 confocal microscope equipped with a 63X oil objective and processed using ImageJ. An *ad hoc* designed macro was used to quantify CD147 signal upon stimulation with EGF[7]. CD147 signal was highlighted applying an intensity-based threshold (Default method), and then fluorescence intensity per field was calculated using the "Measure" command, limiting measurements to threshold. This value was then divided by the number of nuclei in the field, counted using the DAPI signal, to calculate the CD147 fluorescence intensity per cell.
To quantify internalized EGFR/EGF/Tf, cells plated on glass coverslips were stimulated with high dose of different ligands (Alexa-EGF, EGF, TGFα) or Tf at 37 °C for indicated time. Before fixation, samples were subjected to acid wash treatment to visualize only internalized ligand. Different fields of view for each sample were imaged and quantified by Integrated Density (IntDen) to evaluate internalized signal intensity which was divided by the number of cells per field of view. This value was then divided by the number of nuclei in the field and counted using the DAPI signal to calculate the EGF fluorescence intensity per cell.

**Immunofluorescence (IF)**
Cells were plated on glass coverslips in order to have 50% confluency on the day of the experiment. For PLCγ-HA recruitment to PM upon EGF stimulation, cells were serum starved for 2 h prior to stimulation for 2 min with 100 ng/mL of EGF-Alexa-488. Then, cells were transferred on ice, washed with ice-cold PBS and fixed in 4% paraformaldehyde for 10 min,

washed with PBS and permeabilized in 0.1% Triton X-100, 1% BSA/PBS for 8 min at RT. To prevent non-specific binding of the antibodies, cells were then incubated with blocking solution (1% BSA/PBS) for 30 min at RT. Next, cells were incubated for 1 h with primary antibody in blocking solution (anti-HA 1:5000), washed 3 times with 1X PBS and incubated for 30 min with fluorescently labeled secondary antibodies. After 3 washes with PBS, nuclei were DAPI-stained for 5 min and washed again 3 times with 1X PBS.

For Alexa 555-conjugated CTxB staining, cells were stimulated with EGF for 1 min and then fixed in 4% paraformaldehyde for 10 min. Cells were blocked with 1% BSA/PBS for 30 min at room temperature. Before permeabilisation, cells were incubated with 285 ng/mL Alexa 555-conjugated CTxB in 1% BSA/PBS for 30 min at room temperature. Cells were then permeabilised with 0.1% Triton X-100 in 1% BSA/PBS for 8 min at room temperature. After 3 washes with PBS, nuclei were DAPI-stained for 5 min and washed again 3 times with 1X PBS.

Coverslips were mounted with glycerol mounting media. Images were obtained using a Leica TCS SP8 confocal microscope equipped with a 63X oil objective (Leica HCX PL APO CS, 1.4NA) and processed using ImageJ.

**Electron microscopy**

*Pre-embedding immunolabeling.* Serum-starved cells were plated on MatTek 35-mm glass bottom dishes at 70-80% of confluency, incubated with anti-EGFR 13A9 (Genetech) or with anti-CD147 antibody, followed by incubation with rabbit anti-mouse IgG, and, finally, with Protein-A Gold 10 nm (30 min incubation on ice/each step). Cells were then incubated at 37°C for 5 min with 30 ng/mL EGF, as indicated. A control sample left at 37°C for 5 min without EGF, to exclude unspecific antibody internalization. Cells were fixed 1h at RT in formaldehyde/glutaraldehyde 2.5%/1.2% in 0.1M sodium cacodylate buffer pH 7.4 containing 0.5 mg/mL of ruthenium red. After quick washes with 150 mM sodium cacodylate buffer, the samples were post-fixed in 1.3% osmium tetroxide in a 33 mM sodium cacodylate buffer containing 0.5 mg/mL ruthenium red for 2 h at room temperature. Cells were then rinsed with 150 mM sodium cacodylate, washed with distilled water and stained with 0.5% uranyl acetate in $dH_20$ overnight at 4°C in the dark. Finally, samples were rinsed in $dH_2O$, dehydrated with increasing concentrations of ethanol, embedded in Epon and cured in an oven at 60°C for 48 h.

Ultrathin sections (70–90 nm) were obtained using an ultramicrotome (UC7, Leica microsystem, Vienna, Austria), collected, stained with uranyl acetate and Sato's lead solutions, and observed in a Transmission Electron Microscope Talos L120C (FEI, Thermo Fisher Scientific) operating at 120 kV. Images were acquired with a Ceta CCD camera (FEI, Thermo Fisher Scientific).

*Morphometry of NCE TIs and of CCPs.* Morphometry was performed as previously described[7]. Briefly, cellular profiles of thin sections of cells immunolabeled with anti-EGFR antibody (13A9) and stained with ruthenium red were acquired. For the quantification of the number of CCPs or TIs upon different treatment, gold particle clusters present in PM connected (ruthenium red-positive) structures of randomly selected cells were acquired at a nominal microscope magnification of 22000x (pixel size 0.6 nm). Gold clusters were assigned to one of the two categories based on the presence of a clathrin coat and the distance between PM and the tip of the invagination was measured using ImageJ. The number of ruthenium red-positive endocytic structures identified was divided by the PM length measured with ImageJ on acquired low magnification micrographs (nominal microscope magnification of 1200x) and expressed as number of structures per 100 μm of PM.

For quantification of the elongation of endocytic structures the number of short structures were divided by the number of long ones (respectively > 200nm for CCPs and >of 300nm for Tis). Tis structures shorter than 150nm or structures that cannot be unequivocally ascribed to one or the other category were excluded from the analysis.

*KDEL-HRP/DAB visualization of ER.* Cells were fixed in 1% glutaraldehyde in 0.1 M sodium cacodylate buffer (pH 7.4) for 1h at room temperature, followed by incubation with 0.5 mg/mL 3,3'-diaminobenzidine tetrahydrochloride (DAB) and 0.03% hydrogen peroxide in 0.1 M sodium cacodylate buffer pH 6.9 for 20 min at room temperature. Samples were rinsed in sodium cacodylate buffer and post-fixed with 1.5% potassium ferrocyanide, 1% osmium tetroxide in 0.1M sodium cacodylate buffer (pH 6.9) at room temperature. After enbloc staining with 0.5% uranyl acetate in $dH_20$ overnight at 4°C in the dark, samples were dehydrated with increasing concentrations of ethanol, embedded in Epon and cured in an oven at 60°C for 48 h. Ultrathin sections (70 – 90 nm) were obtained using an ultramicrotome (UC7, Leica microsystem, Vienna, Austria), collected, stained with uranyl acetate and Sato's lead solutions, and observed in a Transmission Electron Microscope Talos L120C (FEI, Thermo Fisher Scientific) operating at 120 kV. Images were acquired with a Ceta CCD camera (FEI, Thermo Fisher Scientific).

*ER-PM contact site analysis by EM.* ER-PM contact analysis was performed as previously described[7]. Briefly, For ER labeling, cells were transfected with HRP-KDEL; 24h after transfection, cells were subjected to pre-embedding immunolabeling with anti-CD147 antibodies as described above. Cells were then fixed and processed for EM analysis. To quantify the proximity of ER and CD147, random images of CD147 positive structure were acquired at a nominal magnification of 22K. CD147 positive structures were considered associated or in proximity when at least an ER tubule was present at ≤ 20 nm.

**Measurements of intracellular $Ca^{2+}$ concentration**

*Aequorin measurements.* HeLa cells grown on 13-mm-round glass coverslips at 50% confluence were transfected with the appropriate PM-targeted aequorin chimeras. Aequorin constructs and protocols were previously described[8]. All aequorin measurements were performed in KRB buffer (135 mM NaCl, 5 mM KCl, 0.4 mM $KH_2PO_4$, 1 mM $MgSO_4$, 20 mM HEPES and 5.5 mM glucose, pH 7.4), supplemented with 1 mM $Ca^{2+}$. When EGTA treatment was performed, aequorin measurements were recorded in KRB buffer plus 100 µM EGTA. Stimulation with EGF was performed as specified in the Figure legends. The experiments were terminated by lysing the cells with 0.01% Triton in a hypotonic $Ca^{2+}$-rich solution (10 mM $CaCl_2$ in $H_2O$), thus discharging the remaining aequorin pool. The light signal was collected and calibrated into $[Ca^{2+}]$ values, as previously described[8]. Extent of $Ca^{2+}$ waves was expressed as area under the curve (AUC) and maximal value of peak in box plot graphs (whiskers min to max). Statistical analysis was performed using GraphPad Prism.

*GCaMP measurements* were performed using two different fluorescent $Ca^{2+}$-probes with different affinity for $Ca^{2+}$: GCaMP6f and GCaMP6m[2, 3, 9]. GCaMP6 probes were developed for imaging rapid $Ca^{2+}$ peaks, like the one observed at ER-PM NCE contact sites. To localize the probe to the PM, where NCE contact sites are formed, we added the PM-targeting sequence of SNAP25, as used for Aequorin[7]. GCaMP-stable HeLa clones (PM-GCaMP6f and mitoGCaMP6m) were grown on 35-mm coverslips or MatTek 48 h prior to the acquisition of images. PM-GCaMP6f cells were subjected to 0.5 µg/mL overnight doxycycline induction the day before the recording of the experiment. For image acquisition, cells were washed and kept in KRB buffer supplemented with 1 mM $Ca^{2+}$ and glucose. To determine the PM $Ca^{2+}$ response, cells were placed on a 37˚C thermostat-controlled stage and exposed to 490/406 nm wavelength light using: the Olympus fluorescent microscopy system equipped with a 20x oil objective (Figure 2); Thunder microscope (Leica, DMI8) 40x oil objective (Figure 4); SP8 AOBS (Leica TCS) 20x water objective with 2.5x magnification (Figure 7), acquiring 5fps for a total of 5 minutes. After recording baseline ratio, cells were stimulated with EGF under the indicated conditions. The cells were placed on a 37˚C thermostat-controlled stage and exposed to light of 490/406 nm wavelength. Fluorescent data collected were expressed as emission ratios. The increase in intensity was calculated with ImageJ using an *ad hoc* designed macro and the extent of $Ca^{2+}$ peaks was expressed as AUC and maximal value of peak in box plot graphs (whiskers min to max). Statistical analysis was performed using GraphPad Prism.

**Measurements of mitochondrial membrane potential with TMRM dye**

Cells were grown on 35-mm round glass MatTek at 50% confluence the day before the experiment and then incubated with 1 nM TMRM dye (Catalog Number I34361, T668 Invitrogen) diluted in DMEM without FBS and red phenol supplemented with 2 mM L-glutamine and 25 µM verapamil (V4629, CAS 152-11-4, Sigma Aldrich) for 1 h at 37˚C and 5% $CO_2$. Then, the variation in the intensity of the dye was monitored by time-lapse microscopy. A Leica TCS SP8 AOBS inverted microscope with 20× oil objective was used to take pictures every 2 sec over a 5 min period. The assay was performed using an environmental microscope incubator set to 37°C and 5% $CO_2$ perfusion. After cell incubation with the dye, EGF or TGFα at the indicated doses were added and maintained in the media for the total duration of the time-lapse experiment. To evaluate changes in TMRM levels following stimulation, cells were incubated with a 10 nM diluted solution of the dye in DMEM without red phenol, supplemented with 1% serum, 2 mM L-glutamine, and 25 µM verapamil. EGF of final concentration of 100 ng/mL and remained in the media throughout the entire time-lapse experiment. The same DMEM without red phenol supplemented with 1% serum, 2 mM L-glutamine, and 25 µM verapamil solution was used as the mock control in the absence of EGF. The increase in intensity was calculated with ImageJ using an *ad hoc* designed macro and the extent of the difference in mitochondrial potential was expressed as AUC and maximal value of peak in box plot graphs (whiskers min to max). Statistical analysis was performed using GraphPad Prism.

**Measurements of plasma membrane-endoplasmic reticulum contact sites using the GFP-MAPPER probe**

HeLa cells stably expressing pSLIK-GFP-MAPPER were transfected with siRNAs targeting RTN3 or RAC1, or mock-transfected as a control, as described above. Cells were seeded onto glass coverslips to reach approximately 50% confluency on the day of the experiment. GFP-MAPPER expression was induced by treatment with 1 µg/mL doxycycline for ~16 h before stimulation. Cells were stimulated or not with high-dose Alexa Fluor-labelled EGF (~40 ng/mL) for 5 min, placed on ice, and fixed with 4% paraformaldehyde for 8 min at RT.

To visualize the plasma membrane, cells were stained with an anti-CD147 antibody (BD Biosciences, clone HIM6, #555961) for 30 min at room temperature, followed by an Alexa Fluor 647-conjugated donkey anti-mouse secondary antibody for 30 min. Nuclei were counterstained with Hoechst 33342 (1 µg/mL) for 30 min. All staining procedures were performed without cell permeabilization. Coverslips were mounted using a glycerol-based mounting medium.

Images were acquired using a Leica DMI8 SP8 confocal microscope equipped with a 63× oil-immersion objective. At least 150 cells per condition were analyzed in each independent experiment. GFP-MAPPER signal was quantified using ImageJ by measuring the integrated fluorescence intensity after thresholding and normalizing the values to the number of cells analyzed.

### Super-resolution microscopy

##### Sample preparation

HeLa cells expressing PLCγ-HA were plated on 35 mm dish, No. 1.5 Coverslip, 14 mm Glass Diameter (MatTek). On the following day, cells were serum starved for 2 h and stimulated with 100 ng/mL EGF for 1 min. Immediately after stimulation, cells were transferred on ice, washed with ice-cold PBS and fixed in 4% paraformaldehyde for 10 min, washed with PBS and permeabilized in 0.1% Triton X-100 in 1% BSA/PBS for 8 min at RT. To prevent non-specific binding of the antibodies, cells were then incubated with blocking solution (1% BSA/PBS) for 30 min at RT. After permeabilization, staining was performed with appropriate antibodies against proteins-of-interest (EGFR, HA, RTN3, CLATH) in 1% BSA/PBS, followed by incubation with specific secondary antibodies. Samples were post-fixed in 4% paraformaldehyde.

##### dSTORM imaging

Samples were analyzed by direct stochastic optical reconstruction microscopy (dSTORM) using one of two super-resolution imaging platforms, depending on the experiment. dSTORM is based on the stochastic switching of fluorophores between fluorescent and non-fluorescent states, induced by reducing agents present in the imaging buffer[10]. Maintaining only a sparse subset of fluorescent molecules in each frame allows individual diffraction-limited spots to be spatially separated and their positions to be determined with sub-diffraction precision[11]. Repeated acquisition of the same field and accumulation of the localization coordinates detected in each frame enable the reconstruction of a super-resolution image.

##### Nikon N-STORM acquisition

dSTORM imaging was performed on a Nikon N-STORM microscope configured for oblique-incidence excitation and equipped with an N-STORM module 2 (Nikon Instruments, Tokyo, Japan). Samples were imaged in STORM buffer containing Buffer A, composed of 10 mM Tris, pH 8.0, and 50 mM NaCl, supplemented with the oxygen-scavenging system GLOX. The GLOX system contained 56 mg/mL glucose oxidase (Sigma-Aldrich), 3.4 mg/mL catalase (Sigma-Aldrich), and 1 M mercaptoethylamine (MEA; Sigma-Aldrich).

Alexa Fluor 647 was excited using a 647-nm laser with a nominal power of 120 mW, whereas Cy3 was excited using a 561-nm laser with a nominal power of 70 mW. Both lasers were part of an LU-NV laser unit (Nikon Instruments). For each channel, 15,000 frames were acquired with an exposure time of 20 ms.

For analyses focused on the plasma membrane, only localization events detected at the cell periphery and within 40 nm of the plasma membrane were considered. Nanoscale colocalization was quantified using image cross-correlation spectroscopy (ICCS), as previously described[12]. Briefly, image cross-correlation functions were calculated by averaging the signal over the pixels contained within a selected region of interest. The amplitude parameters obtained from the resulting correlation curves were used to calculate two localization coefficients. Their arithmetic mean was reported as the ICCS colocalizing fraction.

##### Abbelight SAFe MN360 acquisition

Additional dSTORM datasets were acquired using a SAFe MN360 microscope (Abbelight) equipped with two ORCA-Fusion digital cameras (Hamamatsu) and controlled through Abbelight NEO acquisition software.

Samples were imaged in a buffer containing 10 mM Tris-HCl, pH 8.0, 50 mM NaCl, 10% glucose, 0.56 mg/mL glucose oxidase (CZM1045; Cohesion Biosciences), 0.34 mg/mL catalase from bovine liver, approximately 11,000 U/mg (26910.02; Serva), and 50 mM cysteamine (30070; Sigma-Aldrich).
Two-dimensional dual-color single-molecule localization microscopy images were acquired using an UPlanApo 100×/1.5 NA TIRF oil-immersion objective (Olympus) with HiLo illumination. Fluorophores were excited using a 640-nm laser. Dual-color detection was achieved by spectral demixing using a dual-camera detection scheme. For each acquisition, 45,000 frames were recorded with an exposure time of 44 ms.
Single-molecule localization and image reconstruction were performed using Abbelight NEO analysis software. Colocalization analysis was subsequently conducted using Coloc-Tesseler, which generates Voronoi diagrams from single-molecule localization coordinates[13]. Colocalization was quantified using the Spearman rank correlation coefficient.

**Image reconstruction and statistical analysis**

For visualization, super-resolution images from both imaging platforms were reconstructed by representing each detected molecule as a Gaussian distribution centered on its estimated localization coordinates. The width of each Gaussian distribution was determined by the localization precision obtained from the corresponding single-molecule fitting procedure.
At least seven cells were analyzed for each experimental condition. Statistical significance was assessed using two-way ANOVA followed by Šídák's multiple-comparisons test.

**Colocalization analysis by SIM (structured illumination microscopy)**

Hela cells pSlik-GFP-MAPPER were plated on coverslips and GFP-MAPPER was induced by overnight doxycycline (1 µg/mL) incubation. The following day, the cells were serum-starved for 6h and then treated with EGF for 1 min. After stimulation, cells were transferred on ice, washed with ice cold PBS and fixed in 4% PFA for 10 min and subsequently incubated with CtxB (285ng/ml) for 30 minutes at RT. Cells where then permeabilized with Triton 0.1% and nuclei were counterstained with DAPI for 5 min and coverslips were mounted using a glycerol-based mounting medium. Images were acquired using a CrestOptics DeepSIM (Nikon) equipped with an oil-immersion objective lens 60X (1.40 N.A.) and Yokogawa Spinning Disk Field Scanning Confocal System. The analyses were performed through ImageJ software; the percentage of colocalization was calculated using an object-based analysis [JACOP plugin[14]] .

**Bulk RNA sequencing**

Bulk RNA sequencing was performed to assess transcript expression in bulk cell populations. Total RNA was isolated using TRIzol followed by purification with RNeasy Mini columns according to the manufacturer's instructions. RNA quantity was measured by Qubit, and RNA integrity was assessed using a Bioanalyzer, with RIN values ranging from 9.1 to 10. Libraries were prepared from purified RNA using standard RNA-sequencing library preparation methods and sequenced on an Illumina platform according to the manufacturer's instructions. Sequencing was performed to obtain sufficient read depth for transcript detection and downstream analysis. Sequencing reads were processed using standard RNA-seq analysis pipelines and aligned to the human reference genome. Gene expression was quantified to confirm expression of genes of interest in the cell populations. Bulk RNA sequencing was performed for transcript verification and was not used for differential gene expression analysis.

**Mouse study approval**

All mice were maintained in a controlled environment at 18-23°C, 40-60% humidity and with 12 h dark/12 h light cycles in a certified animal facility under the control of the institutional organism for animal welfare and ethical approach to animals in experimental procedures (Cogentech OPBA). All animal studies were conducted with the approval of the Italian Minister of Health (N.1UD) and were performed in accordance with Italian law (D.lgs. 26/2014), which enforces Directive 2010/63/EU of the European Parliament and of the Council of September 22, 2010, on the protection of animals used for scientific purposes.

**Isolation of primary mammary epithelial cells and 3D organoids**

To isolate primary mammary epithelial cells (MECs), a pool of at least two/three FVB mice were sacrificed per genotype and per each biological replicate from 8-15 week-old mice. Inguinal and thoracic mammary glands were excised and subjected to overnight mild enzymatic digestion in medium DMEM-Ham's F12 medium (Gibco, Life Technologies), 20 μg/mL Liberase™ (Roche), 150 U/mL Collagenase Type 3 (Worthington), 1 μg/mL human insulin, 1 μg/mL hydrocortisone (Merck Life science), 10 ng/mL hEGF (Invitrogen), 10 mM HEPES, and 1% Pen/Strep. Samples were incubated at 37°C in a humidified atmosphere containing 5% $CO_2$.

Cell suspension was resuspended in PBS and centrifuged at 335 g for 5 min at room temperature (RT). A three-layer suspension was formed, with the upper layer containing epithelial cells trapped in the fatty layer, a middle layer containing stromal cells that was removed by aspiration and a pellet containing epithelial organoids/MECs and red blood cells. The upper fatty layer and the pellet were harvested and incubated at 37°C with 0.25% Trypsin-EDTA solution for 20 min, pipetting the suspension every 10 min. Trypsin activity was then inhibited with 1X trypsin inhibitor at 1:1 ratio (Merck Life science). If necessary, the suspension was incubated at 37°C with 10 U/mL DNAse I (Merck Life science) for 2 min at RT followed by centrifugation at 335 g for 5 min. The supernatant was removed, and red blood cells were removed with ACK Lysing buffer (Gibco). Samples were then centrifuged at 335 g for 5 min to obtain a pellet of purified MECs. Cells were plated in Mammary Epithelial Cell Growth Medium (MEGM, Lonza) on collagen I-coated plates (Corning® BioCoat™) for 24 h cells at 37°C in a humidified atmosphere containing 5% $CO_2$. 4,000 cells were resuspended in 1:6:3 mix of Rat collagen I (~3-4 mg/mL, Corning): Matrigel (Corning): MEGM in 25 μl drop and plated in Lab-Tek II Glass-Slide (Nunc, 8-Well chambers, 154534). After drop solidification, MEGM medium was added with EGF or TGFα and medium was changed every 2-3 days. Proliferation was monitored after 7 days, with acquisition in brightfield of the entire drop with Thunder microscope (Leica, DMI8, 5X magnification, at different z stacks to cover the entire height of the drop). Quantification was performed on ImageJ, selecting manually each ROI (organoid of interest). Number of organoids and organoid diameters were calculated and reported.

**Isolation of primary intestinal organoids**

The intestines were harvested from 8-15 week-old C57 mice and prepared as described[15]. Briefly, the proximal part of the intestine was collected, opened longitudinally, and washed with ice-cold PBS. The luminal side of the intestine was scraped using a glass slide to remove luminal content and villous structures. After a second wash with ice-cold PBS, the intestine was cut into 2–4-mm pieces with scissors. The pieces were transferred to a tube and further washed with cold PBS (5–10 times) with gentle vortexing. Intestinal fragments were incubated in PBS containing 20 mM EDTA for 20 min on ice. The supernatant was discarded, and cold PBS was added to the fragments. Crypts were released by manually inverting the tube 5–10 times. The supernatant was collected and passed through a 70-μm strainer. The remaining tissue fragments were again resuspended in cold 1% FBS PBS and triturated 5–10 times, and the supernatant was passed through a 70-μm strainer. The previous step was repeated once again. The isolated crypts were then pelleted by centrifugation at 100 × g for 5 min and embedded in Matrigel (Corning). After solidification, growth medium was added and replaced every day. The basal medium consisted of Advanced DMEM/F12 (Thermo Fisher Scientific) supplemented with 1% L-glutamine, 1% penicillin/streptomycin, 1% HEPES, N2 supplement (1×; Thermo Fisher Scientific), B27 supplement (1×; Thermo Fisher Scientific), 1.25 mM N-acetylcysteine (Sigma-Aldrich), 10 mM nicotinamide (Sigma-Aldrich), and gentamicin (100 µg/mL; EuroClone). Growth factors were added as indicated, including murine EGF (0.05 µg/mL; Invitrogen), murine Noggin (0.1 µg/mL; PeproTech), murine R-spondin-1-Fc conditioned medium (in-house; 1:5 dilution), and murine Wnt-3a conditioned medium (in-house; 1:5 dilution). Growth and morphology of organoids were followed using a phase-contrast microscope (Evos) at 2× magnification. Quantification was performed on ImageJ, manually selecting each ROI (organoid of interest). The number of organoids and organoid diameters were calculated and reported.

**Transwell migration assay**

Migration assays were performed using 24-well transwell migration chambers with 8.0 µm pores (Corning®). HeLa cells (30,000/sample) were serum starved for 16h and seeded in the upper chamber of the Transwell insert. The lower chamber contained SS medium as negative control, or SS medium + EGF or TGFα (100ng/mL) as chemoattractant. The Transwell Migration Chambers were incubated at 37°C for 24 h. Then, the upper chamber was cleaned with a cotton swab and cells were fixed with 4% PFA for 10-15 min at RT. Cells were then washed in PBS + glycine 100 mM and stained with Hoechst (1 µg/mL). Migrating cells were acquired with a Nikon Eclipse Ti2 microscope. At least 4 fields of view from each technical replicate were acquired, and two technical replicates were considered in each experiment. Results were reported as migration index (number of cells counted normalized for each independent experiments) counted with ImageJ software (version 2.14.0).

**Statistical analysis and software**

All graphs and statistical analyses were generated using Prism Software V10.2, JMP 19 (SAS) software or Microsoft Excel. All the images were analyzed and quantified with ImageJ 1.54p. Unless otherwise specified, lowercase *n* indicates the number of independent biological replicates, whereas uppercase *N* indicates the number of cells, fields of view, cell tracks as explicitly indicated for each experiment and in the corresponding figure legend. The statistical tests used for each analysis are indicated in the relevant figure legends. Comparisons between two groups were performed using unpaired or paired Student's *t*-tests, as indicated. Multiple-group comparisons were performed using one-way ANOVA. Contingency analyses were performed using Pearson's Chi-squared test. p-value <0.05 *, p-value <0.01 **, p-value <0.005 ***, p-value <0.001 ****.

**References**

1. Mesa, D. *et al.* A tripartite organelle platform links growth factor receptor signaling to mitochondrial metabolism. *Nat Commun* **15**, 5119 (2024).
2. Chen, T.W. *et al.* Ultrasensitive fluorescent proteins for imaging neuronal activity. *Nature* **499**, 295-300 (2013).
3. Patron, M. *et al.* MICU1 and MICU2 finely tune the mitochondrial Ca2+ uniporter by exerting opposite effects on MCU activity. *Mol Cell* **53**, 726-737 (2014).
4. Chang, C.L. *et al.* Feedback regulation of receptor-induced Ca2+ signaling mediated by E-Syt1 and Nir2 at endoplasmic reticulum-plasma membrane junctions. *Cell Rep* **5**, 813-825 (2013).

5. Shin, K.J. *et al.* A single lentiviral vector platform for microRNA-based conditional RNA interference and coordinated transgene expression. *Proc Natl Acad Sci U S A* **103**, 13759-13764 (2006).
6. Uphoff, C.C. & Drexler, H.G. Comparative PCR analysis for detection of mycoplasma infections in continuous cell lines. *In Vitro Cell Dev Biol Anim* **38**, 79-85 (2002).
7. Caldieri, G. *et al.* Reticulon 3-dependent ER-PM contact sites control EGFR nonclathrin endocytosis. *Science* **356**, 617-624 (2017).
8. Bonora, M. *et al.* Subcellular calcium measurements in mammalian cells using jellyfish photoprotein aequorin-based probes. *Nat Protoc* **8**, 2105-2118 (2013).
9. Palamidessi, A. *et al.* Unjamming overcomes kinetic and proliferation arrest in terminally differentiated cells and promotes collective motility of carcinoma. *Nat Mater* **18**, 1252-1263 (2019).
10. Heilemann, M. *et al.* Subdiffraction-resolution fluorescence imaging with conventional fluorescent probes. *Angew Chem Int Ed Engl* **47**, 6172-6176 (2008).
11. Thompson, R.E., Larson, D.R. & Webb, W.W. Precise nanometer localization analysis for individual fluorescent probes. *Biophys J* **82**, 2775-2783 (2002).
12. Pelicci, S. *et al.* Novel Tools to Measure Single Molecules Colocalization in Fluorescence Nanoscopy by Image Cross Correlation Spectroscopy. *Nanomaterials (Basel)* **12** (2022).
13. Levet, F. *et al.* A tessellation-based colocalization analysis approach for single-molecule localization microscopy. *Nat Commun* **10**, 2379 (2019).
14. Bolte, S. & Cordelieres, F.P. A guided tour into subcellular colocalization analysis in light microscopy. *J Microsc* **224**, 213-232 (2006).
15. Nikolaev, M. *et al.* Homeostatic mini-intestines through scaffold-guided organoid morphogenesis. *Nature* **585**, 574-578 (2020).

**Supplementary Figure Legends**

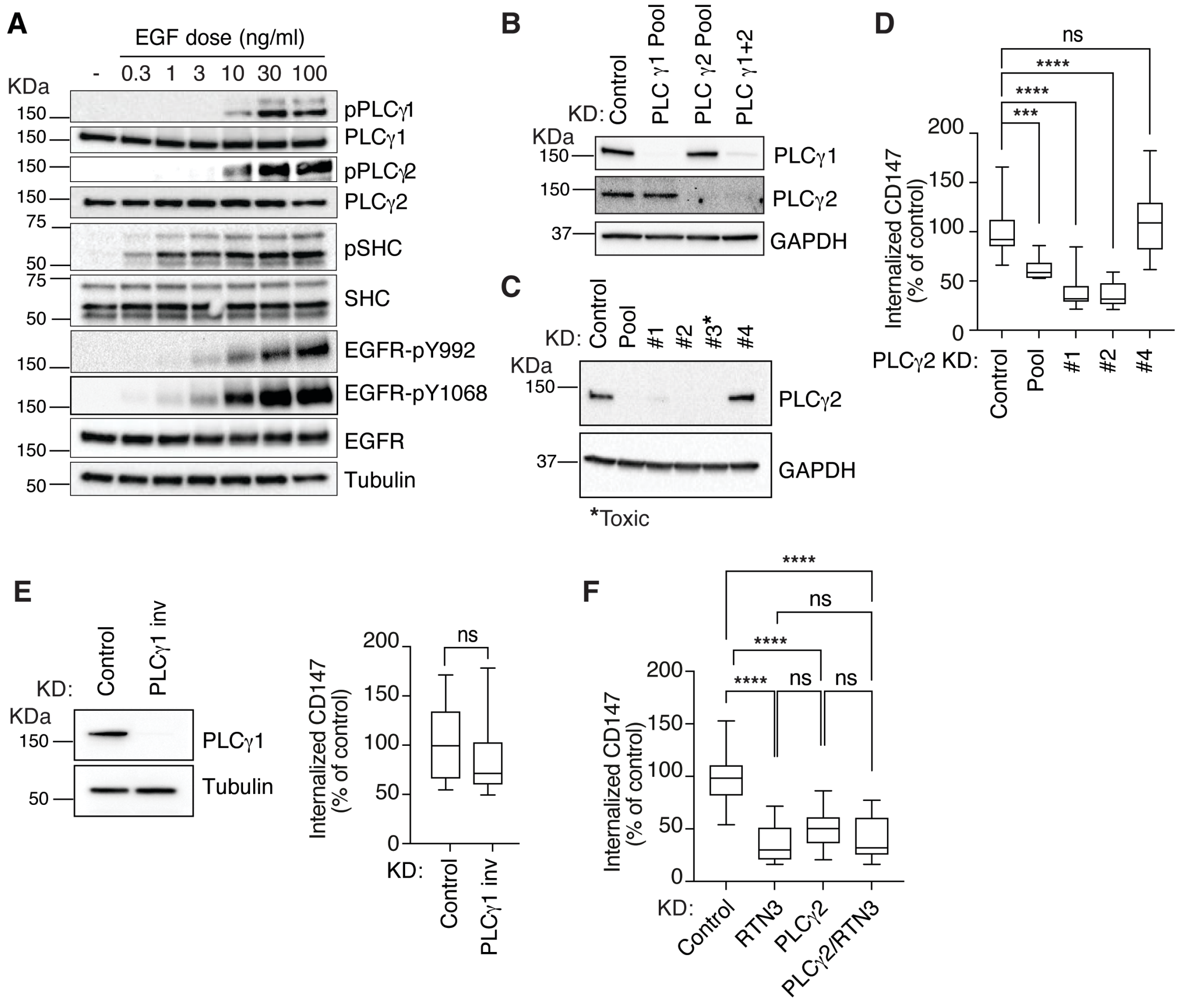


**Figure S1. Additional controls and supporting data for Figure 1. A.** EGF-induced PLCγ phosphorylation correlates with dose-dependent EGFR-NCE activation. HeLa cells were stimulated with the indicated concentrations of EGF for 2 min and harvested for immunoblot analysis of EGFR and downstream signaling. Tubulin, loading control. MW markers shown on the left. Representative experiment shown; n = 2. **B.** PLCγ KD efficiency. HeLa cells were treated with siRNA SMARTpool oligonucleotides targeting PLCγ1 and PLCγ2, individually or in combination. KD efficiency was assessed by immunoblot analysis. GAPDH, loading control. MW markers shown on the left. **C.** Identification of individual siRNA oligonucleotides inducing efficient PLCγ2 KD. The efficacy of the four PLCγ2 siRNA oligonucleotides comprising the SMARTpool was assessed by immunoblot analysis. GAPDH, loading control. MW markers shown on the left. Oligo #2 was chosen for subsequent experiments. Oligo #3 was highly toxic, whereas oligo #4 was ineffective. **D.** Effect of individual PLCγ2 oligonucleotides on EGFR-NCE. CD147 internalization was assessed by IF in HeLa cells treated with the indicated PLCγ2 oligonucleotides or mock control and stimulated with high-dose Alexa Fluor-555 EGF (~30 ng/ml) for 8 min. Quantification of internalized CD147, expressed as percentage of control is shown (box plots of mean integrated fluorescence intensity). N (fields), Control = 10, Pool = 11, Oligo #1 = 11, Oligo #2 = 10, Oligo #4 = 11; n = 1. One-way ANOVA: ***$P < 0.001$; ****$P < 0.0001$; ns. **E.** Effect of an independent PLCγ1 oligonucleotide on EGFR-NCE. Left, KD efficiency of the Invitrogen PLCγ1 oligonucleotide (PLCγ1 inv) was assessed in HeLa cells by immunoblot analysis. Tubulin, loading control. MW markers shown on the left. Right, effect of the independent PLCγ1 oligonucleotide on CD147 internalization was monitored by IF in HeLa cells stimulated with high-dose Alexa

Fluor-555 EGF (~30 ng/ml) for 8 min. Quantification of internalized CD147, expressed as percentage of the mock-treated control is shown (box plots of mean integrated fluorescence intensity). N (fields), Control = 15, PLCγ1 inv KD = 15; n = 1. Student's unpaired t-test; ns. **F.** Depletion of both PLCγ2 and RTN3 does not have additive effects on EGFR-NCE inhibition. CD147 internalization was monitored by IF in HeLa cells subjected to the indicated KDs or mock treatment and stimulated with high-dose Alexa Fluor-555 EGF (~30 ng/ml) for 5 min. Quantification of internalized CD147, expressed as percentage of control is shown (box plots of mean integrated fluorescence intensity). N (fields), Control = 20, RTN3 KD = 20, PLCγ2 KD = 19, RTN3/PLCγ2 KD = 20; n = 2. One-way ANOVA: ****$P < 0.0001$; ns.

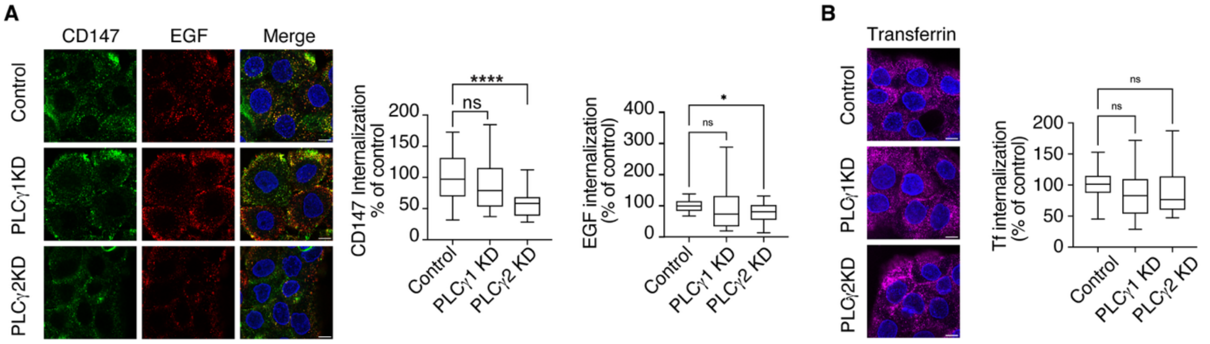


**Figure S2. PLCγ2-dependent EGFR-NCE is conserved in HaCaT cells. A.** CD147 internalization was assessed by IF in HaCaT cells following the indicated KDs or mock treatment and stimulation with high-dose Alexa Fluor-555 EGF (~30 ng/ml) for 8 min. Left, representative confocal images: CD147, green; EGF, red; DAPI, blue. Scale bar, 10 µm. Right, quantification of internalized CD147 and EGF, expressed as percentage of control (box plots of mean integrated fluorescence intensity). CD147: N (fields), Control = 27, PLCγ1 KD = 25, PLCγ2 KD = 23; n = 2. One-way ANOVA: ****$P < 0.0001$; ns. EGF: N (fields), Control = 21, PLCγ1 KD = 24, PLCγ2 KD = 22; n = 2. Welch's t test: *$P < 0.05$; ns. **B.** CME is unaffected by PLCγ depletion in HaCaT cells. Cells were treated as in A, except that Alexa Fluor-488 transferrin (Tf; 50 µg/ml) was used. Left, representative confocal images: Tf, magenta; DAPI, blue. Scale bar, 10 µm. Right, quantification of internalized Tf, expressed as percentage of control (box plots of mean integrated fluorescence intensity). N (fields), Control = 20, PLCγ1 KD = 22, PLCγ2 KD = 21; n = 2. One-way ANOVA: ns.

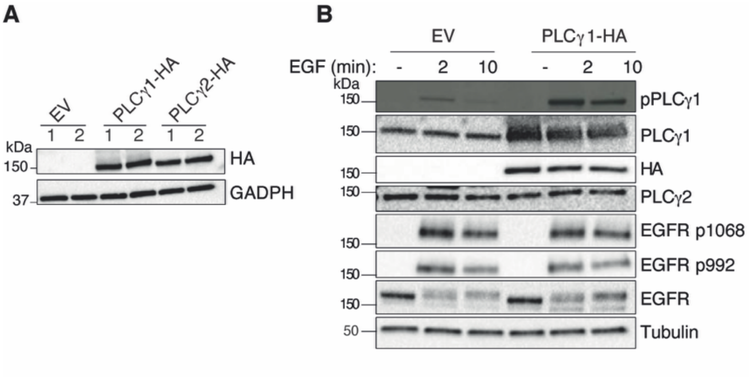


**Figure S3. Additional characterization of stable HeLa cell lines expressing HA-tagged PLCγ. A.** Immunoblot analysis of HA-tagged PLCγ protein expression in stable HeLa cell lines derived from WT HeLa cells (1) or the HeLa doxycycline-inducible clathrin-KD clone 53 (2). EV, empty vector control. GAPDH, loading control. MW markers shown on the left (n = 1). **B, C.** EGFR activation is preserved in stable PLCγ-HA cell lines. Immunoblot analysis of EGFR and PLCγ1 (B) or PLCγ2 (C) activation in stable WT HeLa cell lines stimulated or not (−) with high-dose EGF (100 ng/ml) for the indicated times. Tubulin, loading control. MW markers shown on the left (n = 1).

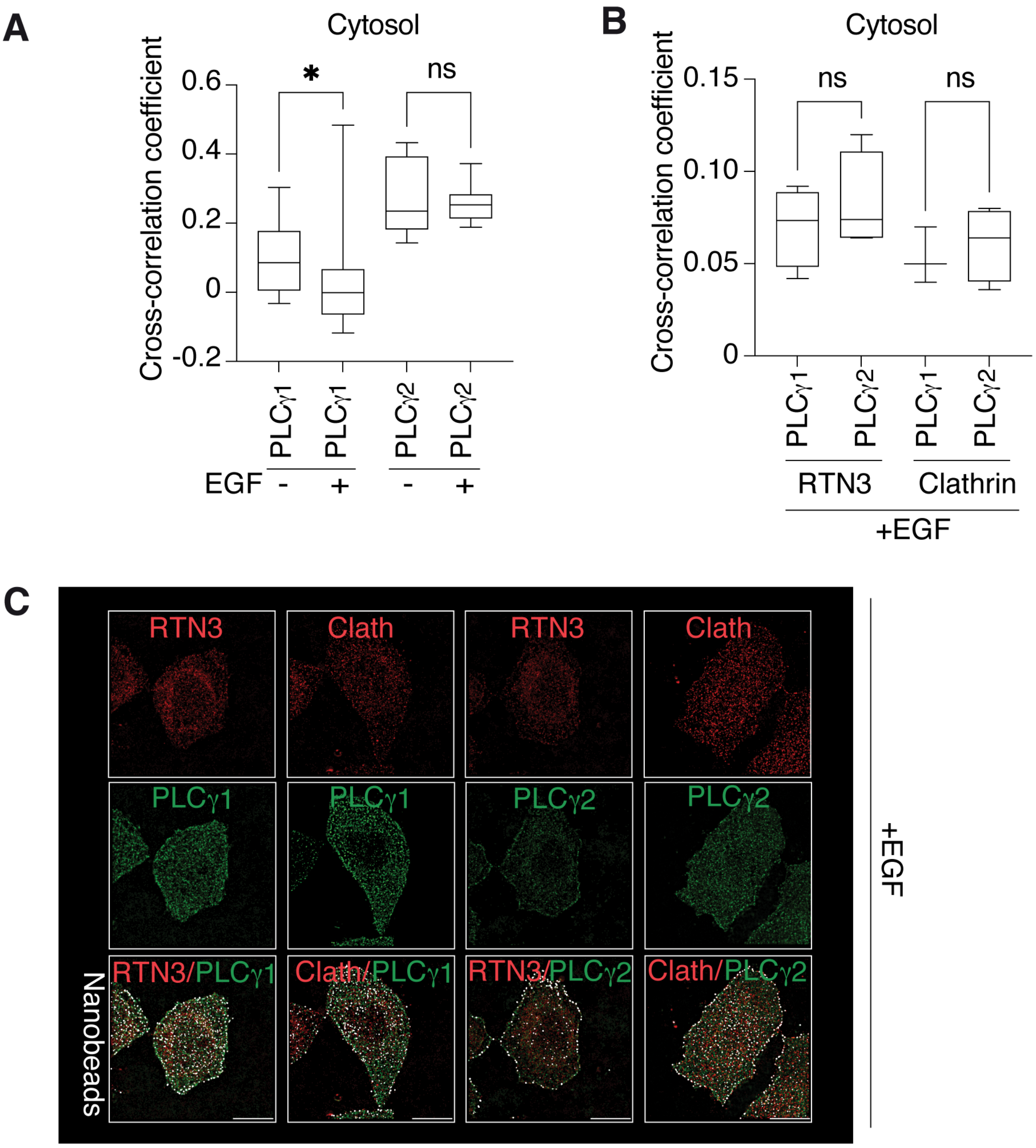


**Figure S4. Additional controls and supporting data for Figure 3. A.** EGF does not induce co-clustering of PLCγ enzymes and EGFR in the cytosol (supporting **Figure 3B**). HeLa cells were stimulated (+) or not (−) with high-dose EGF (100 ng/ml) for 1 min and analyzed by STORM microscopy. Quantification of EGFR and PLCγ-HA co-clustering in cytosol by Spearman cross-correlation analysis, is shown. N (cells): −EGF, PLCγ1 = 12, PLCγ2 = 24; +EGF: PLCγ1 = 11, PLCγ2 = 26. One-way ANOVA, *, $P<0.05$; ns. **B.** PLCγ enzymes exhibit comparable co-clustering with RTN3 or clathrin (Clath) in the cytosol (supporting **Figure 3C**). HeLa cells were stimulated with high-dose EGF (100 ng/ml) for 1 min and analyzed by STORM microscopy using fluorescent nanodiamond registration. Quantification of PLCγ-HA co-clustering with RTN3 or clathrin in cytosol, determined by Spearman cross-correlation analysis, is shown. N (cells): RTN3 PLCγ1 = 4, RTN3 PLCγ2 = 4, Clathrin PLCγ1 = 3, Clathrin PLCγ2 = 4; n = 1. One-way ANOVA, *, $P<0.05$; ns. **C.** Representative STORM images corresponding to the experiment shown in B. Median optical section RTN3/Clath (red), PLCγ-HA (green), and fluorescent nanodiamonds (white) used for image correction and two-channel alignment (same figures as in **Figure 3C**). Scale bar, 10 µm. Channels: RTN3/Clathrin (Channel 1), PLCγ-HA (Channel 2), nanobeads (Channel 3).

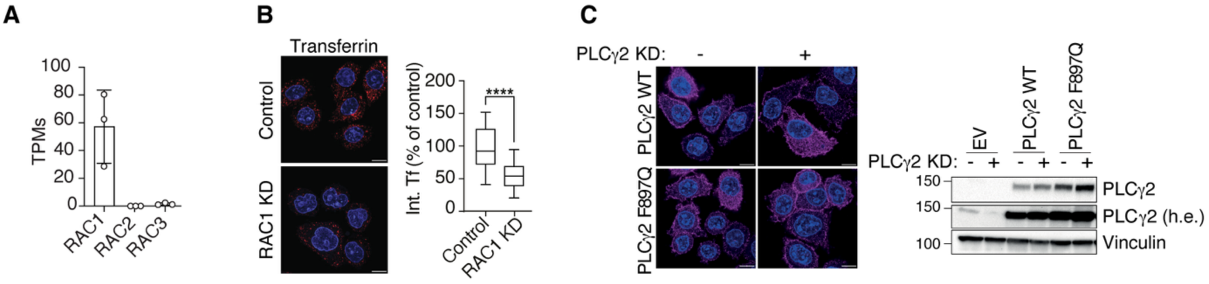


**Figure S5. Additional controls and supporting data for Figures 4 and 5. A.** HeLa cells predominantly express RAC1. mRNA expression levels of RAC1, RAC2 and RAC3 in proliferating HeLa cells, expressed as transcripts per million (TPM; mean ± SD); n = 3. **B.** RAC1 depletion impairs CME. Transferrin (Tf) internalization was monitored by IF in HeLa cells subjected to RAC1 KD or mock treatment and stimulated with Alexa Fluor-555-Tf (50 µg/ml) for 8 min. Left, representative confocal images: Tf, red; DAPI, blue. Scale bar, 10 µm. Right, quantification of internalized Tf (Int. Tf), expressed as percentage of control (mean integrated fluorescence intensity). N (fields): Control = 19, RAC1 KD = 20; n = 2. Student's unpaired t-test; ****, P<0.0001. **C.** Characterization of stable HeLa cell lines expressing siRNA-resistant PLCγ2 constructs. Supporting data for **Figure 5C**. Left, IF analysis of PLCγ2 expression (magenta) in the indicated HeLa cell lines subjected (+) or not (−) to PLCγ2 KD. Scale bar, 10 µm. Right, immunoblot analysis of PLCγ2 expression in the corresponding cell lines. h.e., higher exposure. Vinculin, loading control. MW markers shown on the left.

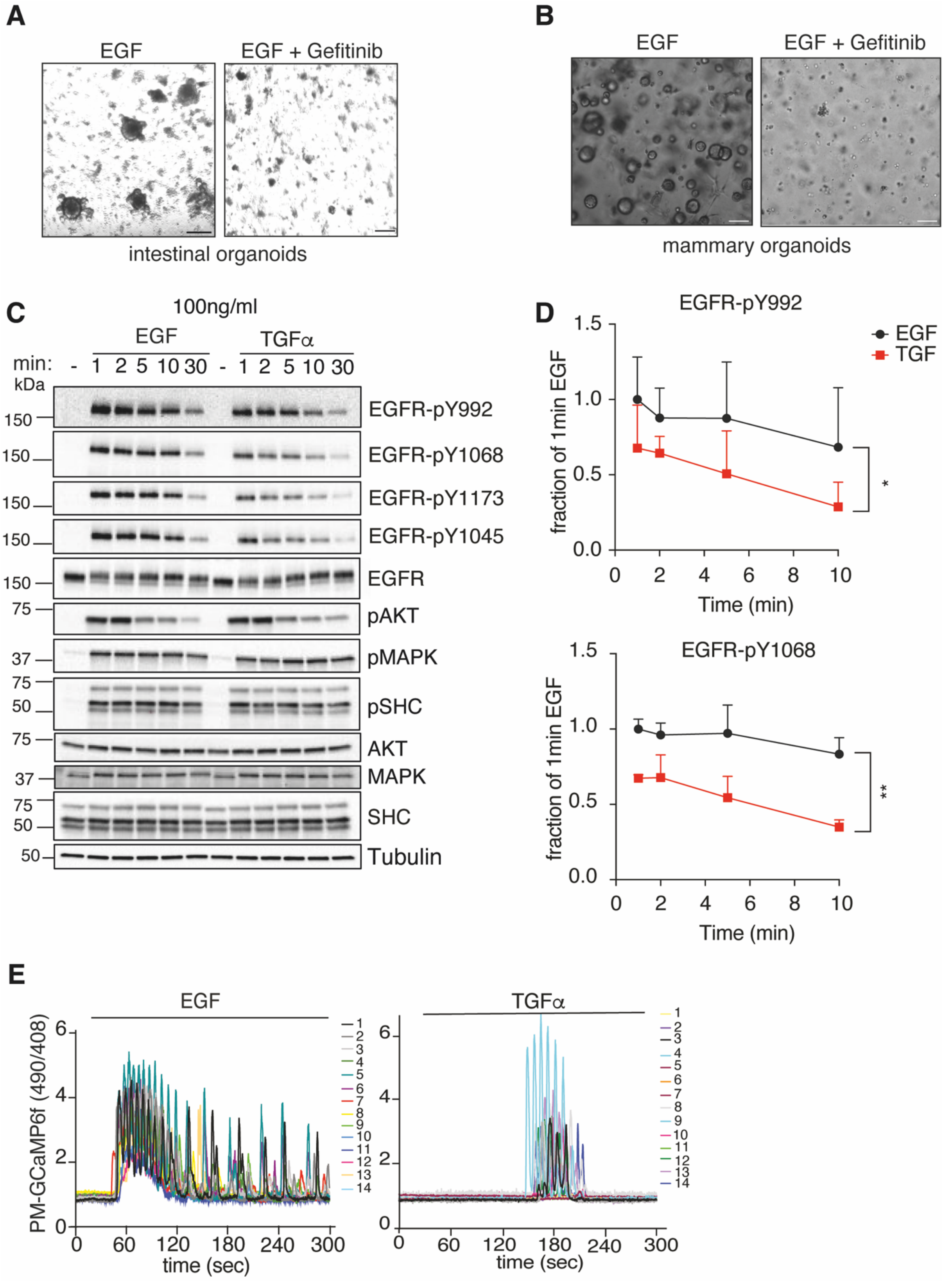


**Figure S6. Additional controls and supporting data for Figures 6 – 8. A.** Intestinal organoid growth requires EGFR signaling. Supporting data for **Figure 6C.** Intestinal organoids derived from FVB WT mice were grown in Matrigel for 7 days in the presence of EGF (50 ng/ml) and the EGFR inhibitor gefitinib (10 μM) or vehicle control. Representative brightfield images of intestinal organoids after 5 days of treatment are shown. Scale bar, 200 μm. **B.** Mammary organoid growth requires EGFR signaling. Supporting data for **Figure 6D.** Mammary organoids derived from FVB WT mice were grown in a Matrigel-Collagen I mix for 8 days in the presence of EGF (50 ng/ml) and gefitinib (10 µM) or vehicle control. Representative brightfield images of mammary organoids after 6 days of treatment are shown. Scale bar, 100 µm. **C,D.** Comparison of EGFR and downstream signaling activation by EGF and TGFα. Supporting data for **Figure 7A.** C, HeLa cells were serum-starved for 16 h, stimulated with high-dose EGF or TGFα (100 ng/ml), and harvested for immunoblot analysis at the indicated times. Representative blot shown (n = 3). Tubulin, loading control. MW markers shown on the left. Note that EGFR-pY992, EGFR-pY1068, and EGFR images are the same as those shown in **Figure 7A**. D, Quantification of EGFR-pY992 and EGFR-pY1068 levels, expressed as a fraction of the EGF-stimulated sample at 1 min (mean±SEM; n = 3). AUC was calculated for the three independent experiments and statistical significance was assessed by the Ratio paired t-test, $^{**}P < 0.01$, $^{*}P < 0.05$. **E.** EGF and TGFα induce distinct PM-localized $Ca^{2+}$ responses. Supporting data for **Figure 8C.** HeLa cells expressing the PM-GCaMP6f $Ca^{2+}$ sensor were stimulated with high-dose EGF or TGFα (100 ng/ml). Multiple $Ca^{2+}$ traces (490/406 nm emission ratio) from individual cells are shown, with each trace represented by a different color.
